# Predictive coding as a unifying principle for explaining a broad range of brightness phenomena

**DOI:** 10.1101/2020.04.23.057620

**Authors:** Alejandro Lerer, Hans Supèr, Matthias S.Keil

## Abstract

The visual system is highly sensitive to spatial context for encoding luminance patterns. Context sensitivity inspired the proposal of many neural mechanisms for explaining the perception of luminance (brightness). Here we propose a novel computational model for estimating the brightness of many visual illusions. We hypothesize that many aspects of brightness can be explained by a predictive coding mechanism, which reduces the redundancy in edge representations on the one hand, while non-redundant activity is enhanced on the other (response equalization). Response equalization is implemented with a dynamic filtering process, which (dynamically) adapts to each input image. Dynamic filtering is applied to the responses of complex cells in order to build a gain control map. The gain control map then acts on simple cell responses before they are used to create a brightness map via activity propagation. Our approach is successful in predicting many challenging visual illusions, including contrast effects, assimilation, and reverse contrast.

**Author summary:** We hardly notice that what we see is often different from the physical world “outside” of the brain. This means that the visual experience that the brain actively constructs may be different from the actual physical properties of objects in the world. In this work, we propose a hypothesis about how the visual system of the brain may construct a representation for achromatic images. Since this process is not unambiguous, sometimes we notice “errors” in our perception, which cause visual illusions. The challenge for theorists, therefore, is to propose computational principles that recreate a large number of visual illusions and to explain why they occur. Notably, our proposed mechanism explains a broader set of visual illusions than any previously published proposal. We achieved this by trying to suppress predictable information. For example, if an image contained repetitive structures, then these structures are predictable and would be suppressed. In this way, non-predictable structures stand out. Predictive coding mechanisms act as early as in the retina (which enhances luminance changes but suppresses uniform regions of luminance), and our computational model holds that this principle also acts at the next stage in the visual system, where representations of perceived luminance (brightness) are created.

## Introduction

Visual perception is relative rather than absolute; the visual system (VS) computes the perceptual attributes of a visual target not only based on its physical properties, but also by considering information from the surrounding region of the target (context). For example, it is possible to induce different kinds of effects by context modification, such that the brightness of a target is contrasted (increasing brightness differences) or assimilated (decreasing brightness differences) with respect to its adjacent surround. Variants of these effects give rise to a myriad of visual illusions, which are of great utility for building hypothesis about computational mechanisms or perceptual rules for brightness perception.

At first sight it seems that contrast effects, such as simultaneous brightness contrast (SBC), can be explained by lateral inhibition between a target (center) and its context (surround). However, activity related to brightness contrast does possibly not occur before V1, albeit the receptive fields of retinal ganglion cells are consistent with lateral inhibition [1].

Unlike contrast, brightness assimilation pulls a target’s brightness towards to that of its immediate context, and therefore cannot be explained by mechanisms based on plain lateral inhibition. In fact, the neural mechanisms involved in generating brightness appear to be more intricate, and only few computational proposals were made so far for explaining assimilation effects. Note that any (computational) proposal could only be deemed as being successful if it explained contrast and assimilation effects at the same time. For example, the simplest (low-level) explanation for assimilation is spatial lowpass filtering, but it would predict brightness assimilation also for contrast displays. Similarly, [2] suggested a retinal model based on a second-order adaptation mechanism induced by double opponent receptive fields; it predicted assimilation, but then again it failed to predict contrast displays. With respect to mid-or higher-level processing, [3] suggested a pattern-specific inhibition mechanism acting in the visual cortex, which inhibits regularly arranged patterns of a visual stimulus.

Anchoring theory is a rule-based mechanism for segregating the contributions of illumination and reflectance in brightness, and thus to map luminance to perceived gray levels [4]. Accordingly, a visual image is first divided into one or more perceptual frameworks (a framework is a set of surfaces that are grouped together). Within each framework, the highest luminance value is anchored at perceived white, and smaller luminance values are mapped to gray levels according to their luminance-ratio with the anchor. Note that “white” may not be the only possible anchor. For example, the lowest luminance may be anchored at “black”, or the mean luminance may be anchored at the Eigengrau level. Although anchoring theory is successful in explaining the perception of reflectance for simple displays, it is not clear how its rules apply to natural images [5, 6]. Furthermore, rather than specifying computational mechanisms, anchoring theory (but also White’s suggestion) represents only a phenomenological proposal, and it is not clear how its empirically defined set of rules could be related to any specific information processing strategy in the visual system (but see e.g., [7]).

Another fairly popular hypothesis holds that contrast and assimilation effects are related to the contrast sensitivity function of the visual system [8]. In one or the other way, this idea forms the base of many computational brightness models, which decompose a visual image according to its spatial frequency content (multi-scale models). Typically, multi-scale models adjust the response amplitudes of spatial-frequency selective filters in a way that the resulting distribution (response amplitude versus spatial frequency) follows the shape of the contrast sensitivity function. Contrast and assimilation effects are then predicted to occur as a consequence of the adjustment.

Specifically, [9, 10] proposed a multi-scale model which represents a luminance image with oriented difference of Gaussians (ODOG) filters. The crucial mechanism in the ODOG-model is (local) contrast normalization for adjusting the spatial frequency representation. In agreement with [8], [9] observed that contrast and assimilation displays are distinguished according to their respective content in spatial frequencies: Luminance patterns with lower spatial frequencies than the encoding filters exhibit a loss of low spatial frequency information in their brightness representation and produce contrast as a consequence. Assimilation, on the other hand, is generated when the high spatial frequencies of a luminance pattern exceed the highest spatial frequencies that can be represented with the encoding filters. Although the ODOG model predicts SBC and White’s effect, it cannot account for more sophisticated visual illusions such as the Dungeon Illusion and the Benary Cross. Both of the latter cannot be reproduced by building a brightness representation with oriented filters that are confined to a specific spatial frequency range. Several extensions of the ODOG model have been proposed in order to account for a bigger set of brightness illusions [11]. However, models based on adjusting the spatial frequency distribution of filter responses fall short of predicting brightness when the input images are contaminated by band-pass filtered noise [12].

Computationally, the spatial operation of certain types of retinal ganglion cells can be dumbed down to taking the second spatial derivative of the visual input. In this way, sharp contrasts (edges or boundaries) are enhanced. Edge integration models [13, 14] aim at inverting spatial differentiation by three successive operations: Contrast extraction (differentiation), thresholding (suppress shallow gradients due to illumination effects), and integration (across edges in order to estimate lightness). Filling-in models [15, 16] therefore may be considered as a neural implementation of edge integration, where boundary activity is propagated laterally in order to generate representations of surface properties. Unlike the spatial filtering models, the integration models therefore encode the input image by their boundaries or contrasts (which may be modulated according to local luminance, cf. [17]). Many filling-in models distinguish two types of boundaries [18–20]. The first type represent barriers for activity propagation, and define the boundary contour system (BCS). The second type is the feature contour system (FCS) and represents surface properties to be filled-in, such as brightness, lightness, color and depth. BCS and FCS were suggested to interact in V4 [21].

The BCS is usually identified with contrast sensitive neurons in the early visual areas (e.g. simple cells and complex cells). BCS activity may be determined with oriented operators at multiple spatial resolutions (scales). In addition to luminance boundaries, the BCS represents the complete 3-D boundary structure of a visual scene, including boundaries from texture and depth. FCS activity was proposed to lie in the blobs (cytochrome-oxidase staining regions) in V1 and the thin stripes in V2 [21].

Whereas the prediction of brightness contrast normally is straightforward with (most) filling-in architectures, assimilation effects remain challenging. A first attempt was made in [18] with a one-dimensional luminance profile. With this simplified configuration, the authors identified mechanisms within the BCS and FCS about how two (non-)adjacent (luminance) regions could influence each other. Specifically, in the BCS, edge activity may be reduced by nearby luminance steps and their polarity. Therefore, these boundaries may not completely block anymore the propagation of activity in the FCS, and may cause FCS activity to fill into non-adjacent surfaces. In this way, one surface may not just be influenced by the brightness of its immediate surrounding region, but even from further away.

With their selective integration model, [22] used occlusion cues for explaining brightness illusions, like White’s effect and the Benary Cross. The trigger cue for separating boundaries in depth are T-junctions, which are nevertheless effective in flat luminance displays, too. In their model, the original boundary structure (BCS) is modified such that boundaries along the stem of the “T” are suppressed, while the others are enhanced. The modified boundary map (“context boundaries”) is subsequently used for suppressing (in the FCS) contrast measurements at spatially corresponding positions. A filling-in process then uses the original boundaries along with the modified contrast measurements. However, psychophysical evidence suggests that White’s illusion seems not to be affected significantly if the T-junctions are suppressed [23, 24], nor seem to be other illusions [25]. Furthermore, it is not readily clear whether junction rules represent reliable cues in complex natural scenes: The utility of junctions rules has only been illustrated with relatively simple artificial displays [22, 26].

Another filling-in-like model used center-surround receptive fields at four resolution levels for edge (or contrast) extraction [2]. At each resolution level, filter response amplitudes (“local contrast”) were gain-controlled with a low-pass filtered version of themselves (“remote contrast”). A brightness map was estimated from the gain-controlled contrast map with fixpoint iteration of a Laplacian [27], what implements the filling-in process. The model was successful in simulating assimilation and reverse assimilation effects (mostly centered on challenging variants of White’s effect), but failed in predicting Simultaneous Brightness Contrast (SBC).

More recently, [28] extended the mechanism of [18] for explaining assimilation effects to two dimensions. Thus, Domijan modified BCS activity accordingly before filling-in occurs in the FCS. To this end, he relies heavily on the max-operator, which he justified with a dendritic circuit proposal. In comparison, [29] used a modified diffusion operator, which could be efficiently implemented (both physiologically and computationally) with rectifying gap-junctions or rectifying dendro-dendritic connections [17, 29]. For filling-in, Domijan’s used a luminance sensitive pathway. Luminance-modulated contrast responses are computed with an unbalanced center-surround kernel, similar to [30], but see also [17] for a different way of computing multiplexed contrasts). In addition, [28] computed BCS activity by first deriving a local boundary map, where the loss of activity at junctions and corners was corrected. Based on the local boundary map, a global boundary map was computed. In the latter, contours which are parallel or co-linear to another contour were enhanced. Finally, local boundary activity was divided, at each position, by global boundary activity. The division is approximately one at those positions where no contour enhancement took place in the global boundary map. Otherwise it is smaller than one. The final BCS output keeps only those activities that are relatively close to one – low contrast boundaries which are parallel to high contrast edges are eliminated. This causes FCS activity to freely diffuse across the eliminated boundaries. In this way assimilation displays are predicted. Unfortunately, Domijan did not show any results with real-world luminance images as input, and thus it is not clear whether his proposed mechanisms are robust and how well they generalize.

A completely different approach for explaining visual illusions is based on a statistical analysis of real-world images. This approach suggests that the perception of brightness [31, 32] or lightness [32–34] is related to knowledge about the statistical relationships between visual patterns. In particular, [31] proposed that the brightness of a visual target embedded in some context depends on the expected luminance according to a probability distribution function. The probability distribution function integrates all contexts in that the target was seen previously. The perception of the target then depends on its expected luminance given its current context: It is perceived as brighter if the expected luminance is lower, and it is perceived darker otherwise. This approach is successful in predicting contrast and assimilation for several visual illusions, and suggests a statistical relationship between luminance patterns and brightness perception. Unfortunately, no attempt has been made in order to unveil any information processing strategy from the statistical analysis (but see [34]).

As with some of the models reviewed above, our approach also emphasizes the importance of boundaries in brightness perception. Specifically, we propose to reduce redundancy in the boundary maps. Such encoding strategies usually reduce the overall activity of a representation (and thus the expenditure of metabolic energy, [35, 36], and are also known as efficient coding [37], predictive coding [38], whitening [90] or response equalization [40].

The underlying idea of our computational model is to adjust a boundary map, such that redundant activity is suppressed, while non-redundant activity is enhanced. Since neurons that encode redundant patterns tend to be over-represented, the overall boundary activity is reduced after the adjustment (response equalization). Response equalization is carried out by a dynamic filter. Fig 1 shows an overview of our model. In the first step an input image is encoded in parallel by two sets of Gabor filters, which mimic the spatial response properties of simple cells in V1 [46]. The responses of the high-resolution filters define the contrast-only channel, while responses of the more coarse-grained filters define the contrast-luminance channel. From the contrast-only channel, we compute boundary activity via local energy [42, 43]. Local energy is insensitive to the phase of simple cells, and thus resembles complex cell responses. We learn a decorrelation kernel from the local energy map, and apply it to the latter in order to reduce its redundancy (=dynamic filtering). The redundancy-reduced energy map then functions as a gain control map for both contrast channels. As a consequence, contrast activity is modified. Subsequently, an iterative procedure is used to recover a brightness map from the contrast channels. Our iterative procedure resembles a filling-in process. The utility of our model is demonstrated by predicting many challenging visual illusion, including some that were never predicted by any other computational model so far.

**Fig 1.**
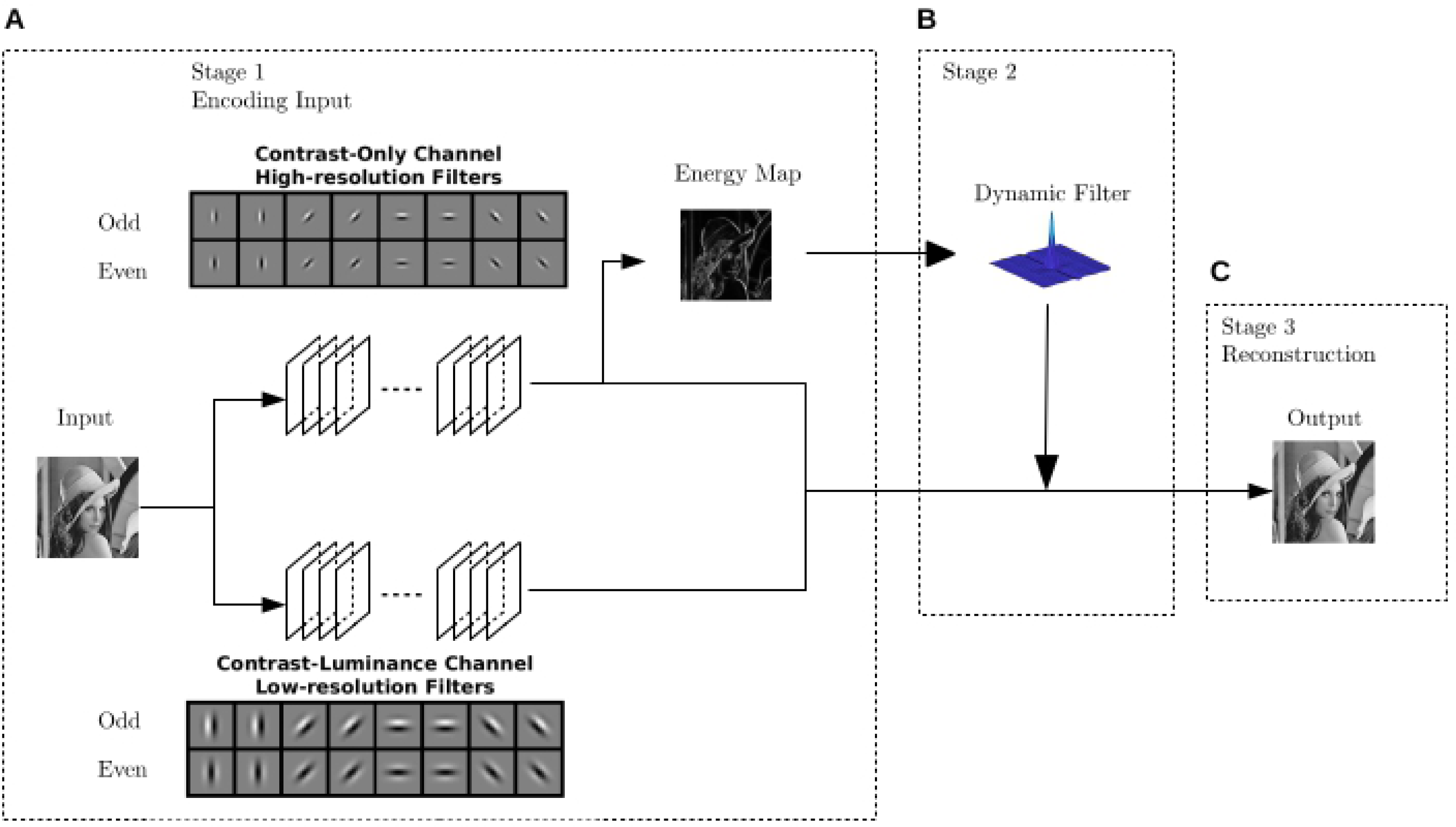
Model overview. Each of the three stages is mathematically specified in corresponding subsection in the text. (A) Stage 1: The Contrast-Only channel and Contrast-Luminance channel are encoded by filtering the input image with a corresponding set of Gabor filters with high spatial resolution and coarse resolution, respectively. The local energy map is computed from the contrast-only channel. (B) Stage 2: The kernel of the dynamic filter is estimated from the local energy map. Dynamic filtering equalizes the amplitude spectrum of the energy map, reducing redundancy. The decorrelated energy map serves as gain control map for both contrast channels. (C) Stage 3: The output of the model is a brightness map that is obtained by solving an inverse problem, that is recovering the image from both of the contrast channels. Thus, the contrast channels do not interact before Stage 3.

## Materials and methods

Fig 1 depicts the three stages of our model. Stage 1 encodes the input image into a contrast-only and a contrast-luminance channel by two respective set of Gabor filters. Stage 2 implements the dynamic filtering process. It consists of equalizing response amplitudes of the local energy map and modifying both contrast channels. Finally, Stage 3 refers to the filling-in process for estimating a brightness map. The brightness map represents the output of our model. The three stages are detailed in the following subsections.

### Stage 1: Encoding

#### Contrast-only and contrast-luminance channel

We use Gabor filters in order to encode contrast-only and contrast-luminance information. In the primary visual cortex, simple cells respond to oriented light-dark bars across a certain spatial frequency range [44] and their receptive fields can be modeled by Gabor filters [45–47]. Consistent with the properties of Gabor filters, it seems that many simple cells in V1 encode contrast information. Under certain circumstances though, neurons in V1 may respond to surface brightness as well, even without (sharp luminance-)contrasts in their receptive fields [48, 49]. For example, [50](found such neurons in V1 which have large receptive fields, broad orientation tuning, and a preference for low spatial frequencies. These neurons respond to both contrast and luminance. Accordingly, we computed the contrast-only channel by Gabor filters with high spatial frequency tuning and balanced ON-OFF sub-regions (i.e., the sum across the kernel is zero), such that they did not respond to homogeneous regions of the stimulus. On the other hand, the contrast-luminance channel is computed by Gabor filters having a lower spatial frequency, and unbalanced ON-OFF sub-regions (i.e., the sum across the kernel is positive) such that they respond to both luminance and contrast. Fig 1 illustrates these two sets of filters (see S1 Appendix for parameter values and a mathematical description). The contrast-only and contrast-luminance channel were computed by simply convolving (symbol “*“) a luminance image with the corresponding set of Gabor filters. That is, if “g” represents a Gabor kernel, then

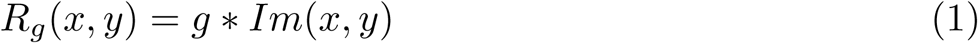

represents its activity. The arguments (x,y) indicate 2D spatial coordinates. The contrast channels remain separated until the filling-in process.

#### Local Energy Map

The idea underlying the “energy map” is to generalize across contrast polarity, orientation and phase, respectively, what also leads to a certain degree of position invariance [42, 43, 51, 52]. The local energy map therefore resembles the properties of complex cells in the primary visual cortex [53]. Particularly, complex cell responses are similar to those of simple cells in terms of orientation and spatial frequency preference, respectively. However, complex cell responses tend to be non-linear and shift-invariant with respect to contrast phase [53, 54]. Local energy is computed from a pair of Gabor filters with quadrature phase (and identical orientation and spatial frequency) by summing their squared responses Fig 2. Finally, the energy map E was computed through averaging the activity of our model complex cells across all orientations. Parameters and mathematical details can be found in S2 Appendix.

**Fig 2.**
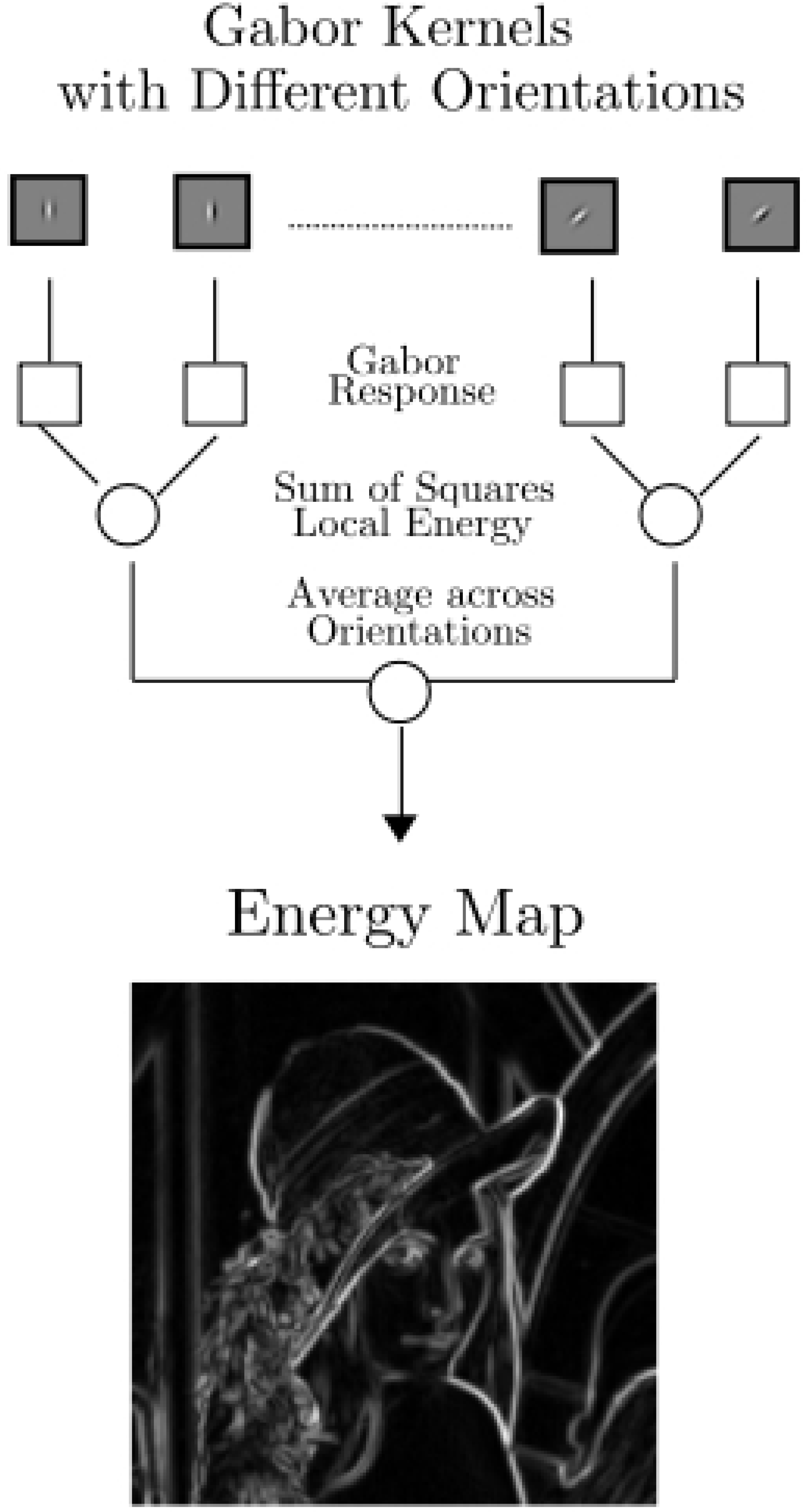
Computation of the energy map. At first, the local energy was computed (at each position in the image) by summing the squared responses of a pair of Gabor kernels which have the same spatial frequency and orientation whereas their spatial phase differ in 90 degrees. After that, an energy map was computed by averaging the activity of local energy across orientations.

### Stage 2: Decorrelation

In this section we describe stage 2 of our model (Figure 1B). The first subsection describes how the dynamic filter is computed using zero-phase whitening (ZCA: [55]. With the dynamic filter we equalize the amplitude spectrum of the energy map. In fact, it produces very similar results to the “Whitening-by-Diffusion” method proposed in [40]. In the second and third subsections we detail the computation of the gain control map and how it interacts with the whitened energy map, respectively.

#### Dynamic Filter

The purpose of the dynamic filter is to equalize the amplitude spectrum of the energy map. We compute our “dynamic filter” using zero-phase whitening (ZCA), a technique which has been used for learning the receptive fields of retinal ganglion cells [55]. ZCA is very similar to principal component analysis (PCA), and signal decorrelation can be achieved with both of the latter. However, the components are constrained to be symmetrical with ZCA. This “symmetry constraint” guarantees that the principal components are localized in the spatial domain [55], and therefore can be used as filter kernels. We nevertheless introduced a couple of modifications to the original ZCA (see S3 Appendix). As a result of the modifications, we obtained a spatial filter that adapts to the spatial structure of the local energy map of an image. It is called “dynamical” because a different filter is learned from each image. After filtering, the amplitude spectrum of the energy map is more uniform (see Fig 3). By the Wiener–Khinchin theorem, a more uniform power or amplitude spectrum implies that the original signal is more decorrelated [56, 57]. For the decorrelated energy map, this means that spatial patterns with low redundancy tend to be intensified, while patterns with high redundancy tend to be attenuated. This is illustrated with Fig 3, where after filtering horizontal edges are intensified in the energy map as compared to vertical ones.

**Fig 3.**
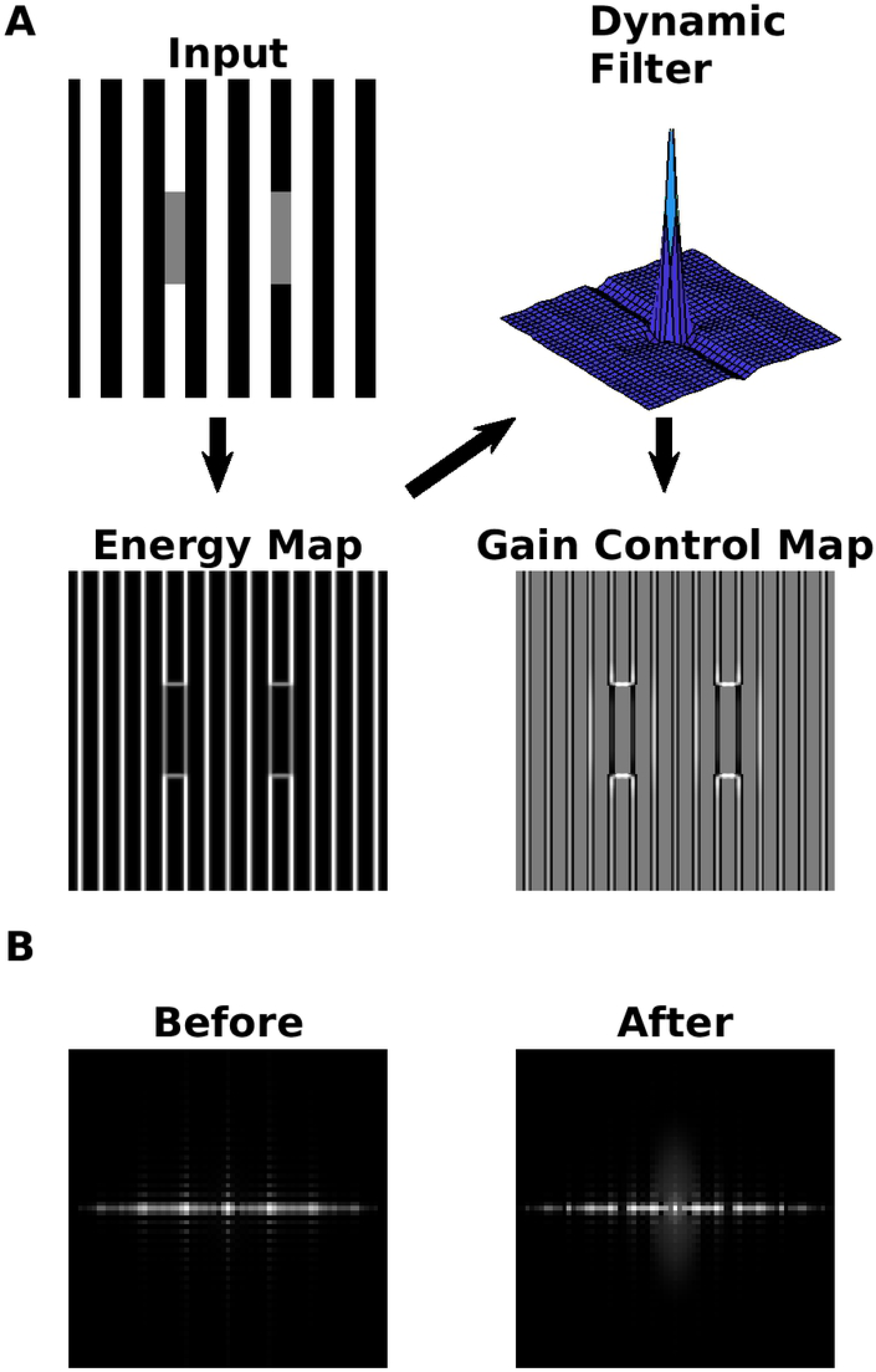
Illustration of the dynamic filter. (A) The arrows indicate the steps in order to obtain the gain control map: (i) A local energy map is computed; (ii) a dynamical filter is constructed with a customized zero-phase whitening procedure (ZCA, see S3 Appendix); (iii) a gain control map is obtained by filtering the energy map with the dynamical filter (see subsection Gain Control Map). (B) The power spectrum (= square of amplitude spectrum) before and after of applying the dynamic filter on the local energy map.

#### Gain Control Map

A gain control map G was computed in two steps. First, the dynamic filter F was used as a convolution kernel for the energy map E:

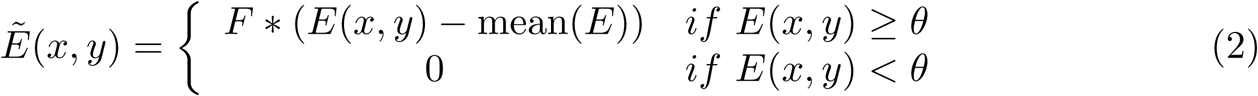

where the symbol “*” indicates convolution. We set the threshold to 10 percent of the maximum activity as *θ* = 0.1max(*E*). In the second step we normalized the activity of the gain control map with a sigmoidal function *S*(*x, a, b*) = 1*/*(1 + *e*^−*ax*−*b*^) as

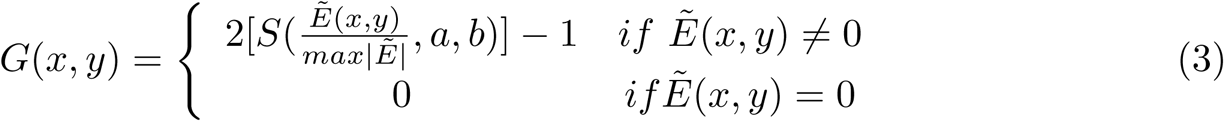

with *a* = 5 and 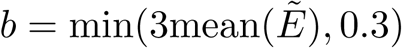. Notice that the gain control map *G* is normalized to [−1, 1]. Fig 3A shows an example of a gain control map.

#### Gain control and normalization between channels

The contrast-only and the contrast-luminance channel were subjected to gain control using the decorrelated energy map as

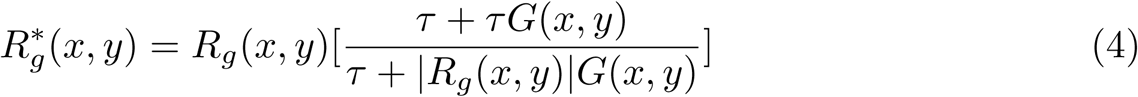

where *G* is the gain control map, *τ* is a control parameter, *R*_*g*_ represents the activity of a Gabor filter *g* of the corresponding filter set, and (*x, y*) are the pixel coordinates of the input image. The value of acts as an upper bound to the maximum activity that we observed during the simulations. If the value of *G* at pixels coordinate (*x, y*) is bigger (or smaller) than zero, then the activity of the corresponding Gabor response will increase (or decrease). On the other hand, if *G*(*x, y*) ≈ 0 then 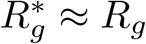 at position (*x, y*). Finally, after applying Eq 4 to each channel, the results were lowpass filtered with a Gaussian kernel (standard deviation 1 pixel) in order to reduce possible artifacts.

### Stage 3: Brightness estimation as a filling in process

The brightness map 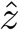 is the output of our model. It is estimated by minimizing an objective function *E*(*z*), which optimizes the trade-off between the reconstruction error (first term in the sum of Eq 5) using the gain-controlled contrast channels and a smoothness constraint (second term):

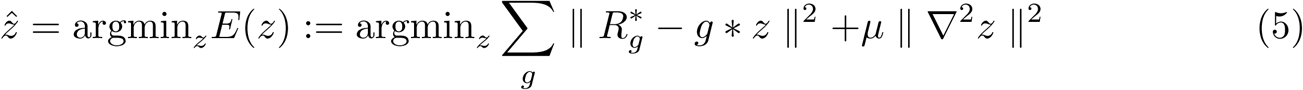

Notice that the sum involves the Gabor filters *g* of both contrast channels (i.e., contrast-only and contrast-luminance). The regularization parameter controls the smoothness constraint and is set to 0.01. The Laplacian is denoted by ∇^2^. The smoothness term serves to reduce artifacts produced at discontinuities. The Eq 5 is solved iteratively with the conjugate gradient method (see S4 Appendix for more detail), which was terminated either when having reached a maximum number of iterations, or when an error criterion was satisfied:

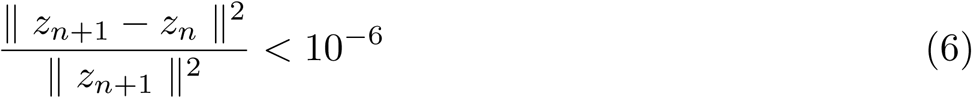

The last formula measures the difference in reconstruction between subsequent iterations (“relative error”). In terms of convergence, we observed as our model converged (globally) to a stable solution in all considered cases.

Our model functions as an edge-integration model, in which the activity is propagated across edges into a certain direction and orientation. Direction and orientation is determined by the odd-symmetrical Gabor filters. Thus, image reconstruction (i.e., brightness estimation) by means of Eq 5 proceeds according to a filling-in process if derivative (i.e., odd Gabor) filters are involved [58]. It is interesting to highlight the role of both channels for estimating brightness. Almost all activity propagation depends on the Contrast-Luminance channel, because its larger filter kernels propagate the activity faster than the narrower filters of the Contrast-only channel. In addition, the model cells of the Contrast-Luminance channel which encode luminance lead to a further reduction of the number of iterations to satisfy the stop criterion. Fig 4 compares the reconstruction of a luminance staircase at 10 iterations if only one of the channels was used. Note, although both channels contribute to Eq 5, brightness estimation depends on the Contrast-Luminance channel in the first place, while the Contrast-only channel serves to encode edge-information.

**Fig 4.**
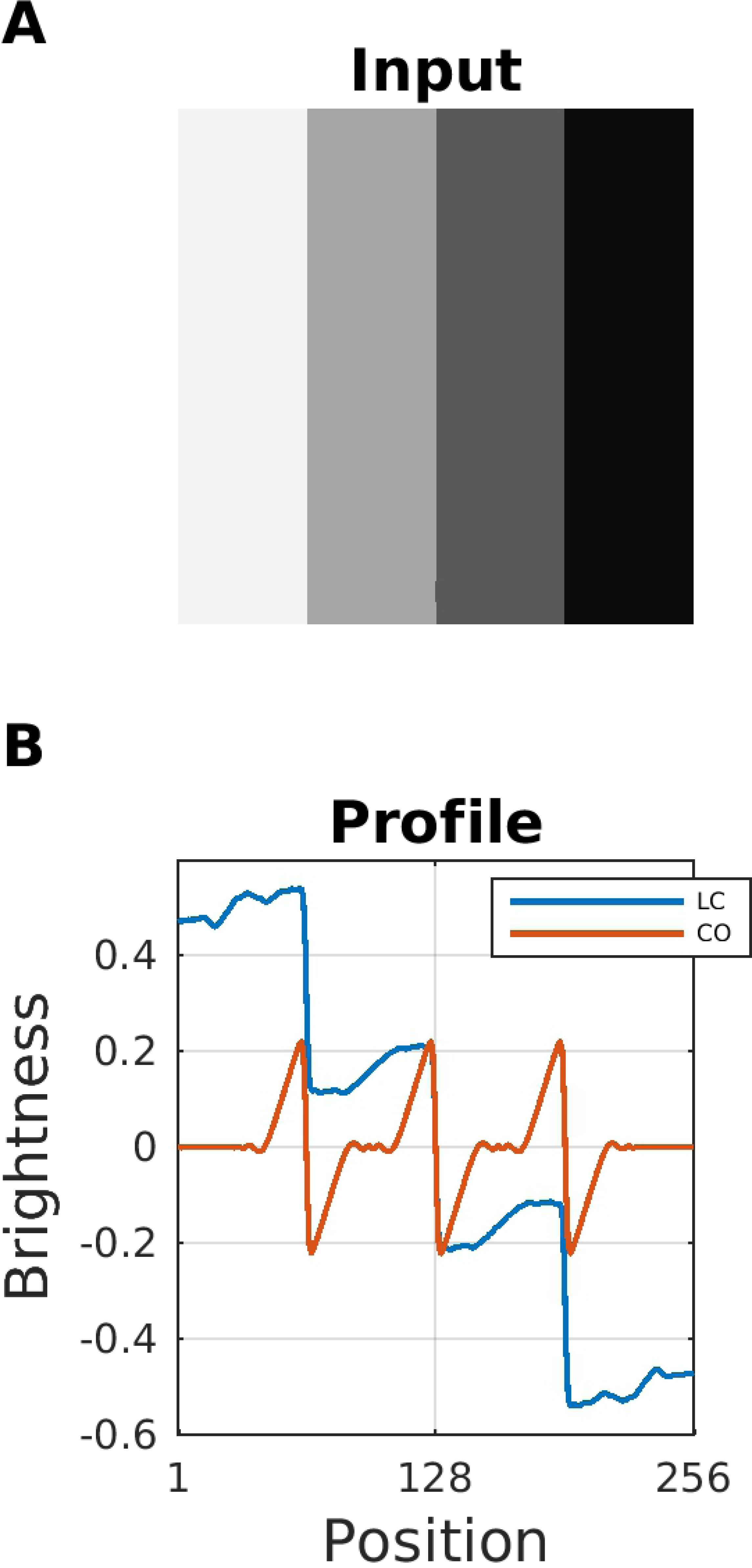
Brightness estimation using different channels. (A) A luminance staircase (giving rise to Chevreul’s Illusion) served as input. (B) The resulting brightness profiles are estimated at 10 iterations. Blue curve (legend label “CL”) uses only the Contrast-Luminance channel in Eq 5. Red curve (“CO”) uses only the Contrast-Only channel but removing responses to luminance.

### Classification of Model Predictions According to Three Scenarios

The effect of the Gain Control Map on estimated brightness (=model output) is as follows. If a Gabor response amplitude after gain control has increased (decreased), it would produce a major (minor) contrast in the reconstructed image (=estimated brightness). This means that brightness estimates as generated by our model depend critically on Eq 4. The numerator depends directly on the Gain Control Map (which in turn is generated by the dynamic filter). The denominator of Eq 4 implements a normalization mechanism. The purpose of this section is to evaluate the relative influence of the numerator and the denominator, respectively, on predicted brightness. To this end, we identified three prominent scenarios for explaining corresponding classes of brightness illusions (Fig 5).

**Fig 5.**
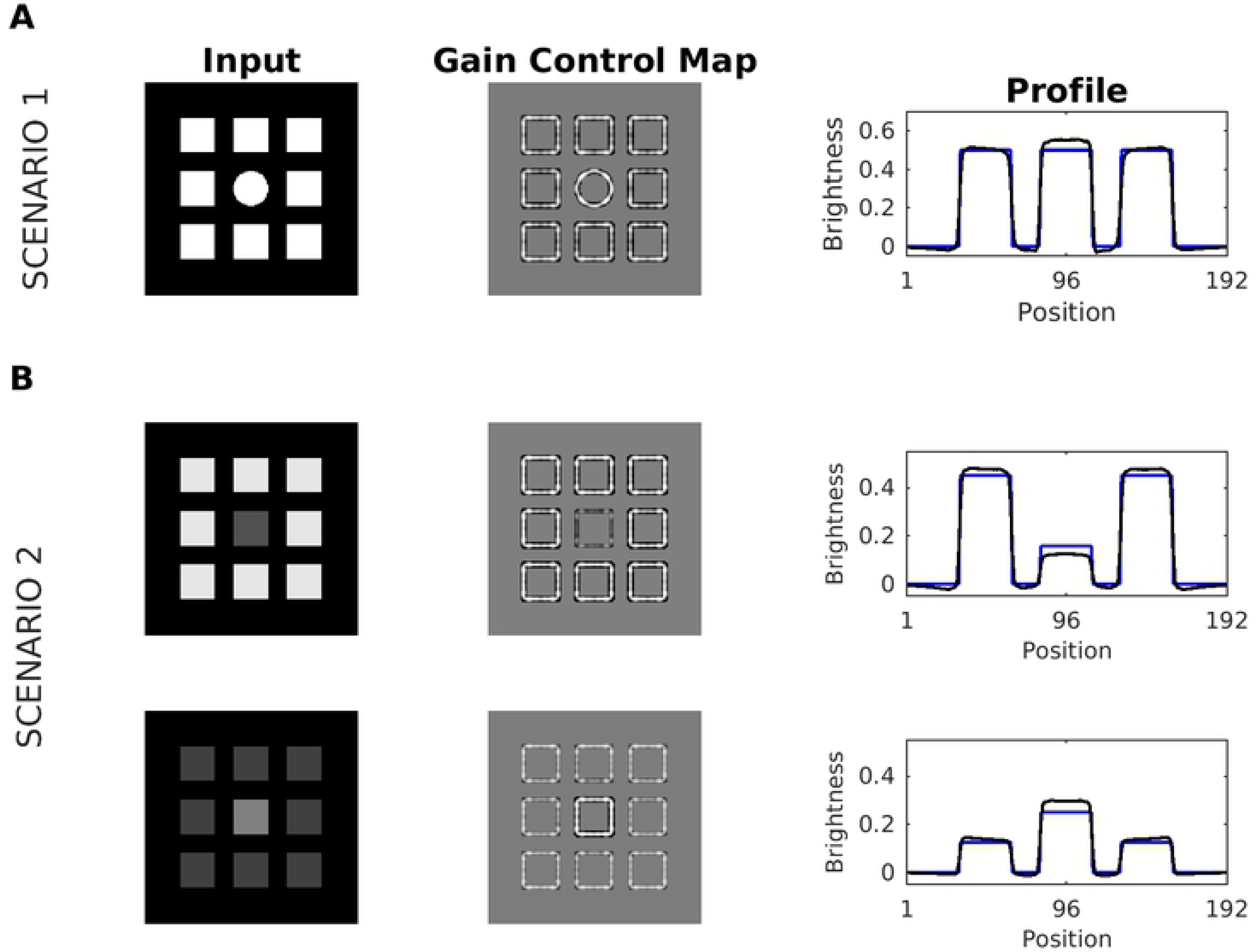
Scenario 1 and Scenario 2. (A) Scenario 1: A disk embedded in a redundant pattern of eight squares served as input (first column). The middle column depicts the corresponding gain control map G, and the right column the profiles of input (blue line) and brightness estimation (black line). The brightness of the center disk ts enhanced with respect to luminance, meaning that a brightness contrast effect is predicted merely based on redundancy (but not on grounds of luminance - note that all features have the same luminance). (B) Scenario 2. The input consists of a series of nine squares arranged in a spatially redundant pattern, where the middle square has a different luminance. The profile plot suggests an overall increase in brightness contrast: Brightness of the middle square is further reduced, while the brightness of the surrounding squares is enhanced. While the brightness contrast also increased in the display with the bright middle square, this increase in contrast is caused nearly exclusively by the middle square.

The luminance pattern giving rise to Scenario 1 differs only in its spatial structure (Fig 5A), as all structures have the same intensity value. In this case, the major contribution to predicted brightness comes from the Gain Control Map (numerator of Eq4). The patterns with high spatial correlations are attenuated by the dynamic filter, while patterns with lower spatial correlation are somewhat increased such that a brightness contrast effect in predicted for the central disk. Scenario 2 is defined by luminance patterns with similar spatial structure but different intensity range (see Fig5B). Analogously to Scenario 1, the major contribution to predicted brightness stems from the Gain Control Map, but here the effect is limited by the size of the dynamic filter. We observed that the dynamic filter will not only reduce the spatial correlations, but it will also act as a contrast filter, if the redundant activity is in a sufficiently small spatial region. As a result, redundant activity with higher (lower) intensity than the other patterns would be increased (decreased). If this increment (or decrement) is sufficiently big, it will produce a major (minor) brightness contrast effect.

In Scenario 3, the major contribution to predicted brightness is caused by the Contrast-Luminance channel and normalization (denominator of Eq 4), while the contribution of the gain control map cancels. Edges in the Contrast-Luminance channel might be enhanced via the Gain Control Map. These enhanced edge positions eventually produce a boost in (estimated) brightness contrast. The degree of boosting depends on the ratio between the activity (after boosting) and the control parameter (upper bound) in Eq 4). An example of this effect can be observed comparing both displays and profiles (at the edges) in FigEq 6) (see description). It is essential to highlight that Scenario 3 serves just to explain Chevreul’s illusion (see results section), but is not relevant to all other brightness displays.

## Results

This section presents simulation results of our model in predicting brightness illusions. The first subsection focuses on contrast effect: Simultaneous Brightness Contrast, Benary-Cross and Reverse Contrast. The second subsection is dedicated to assimilation effects: White effect, Todorović’s Illusion, Dungeon illusion, Checkerboard illusion, and Shevell’s Rings. Finally, the last subsection shows our brightness predictions for the Craik-O’Brien-Cornsweet effect, Hermann/Hering grid, Chevreul’s illusion (including the luminance pyramid), Mach Bands, and Grating induction. It is essential to highlight that in this section, all input images were normalized, so that pixel intensity ranged between −0.5 and 0.5 (unless stated otherwise).

### Brightness Contrast Effects

#### Simultaneous Brightness Contrast (SBC)

The SBC display consists of two gray patches with identical luminance which are embedded in a dark and bright background, respectively. The patch on the bright background is perceived as darker than the patch on the dark background (Fig 7A). SBC can be attributed to low-level processing. For example, retinal ganglion cells may enhance patch contrast by lateral inhibition. However, other studies suggest that SBC may involve higher-level processing as well [59]: The apparent brightness of the patches can be modulated by the region surrounding the patches (=spatial context). In fact, psychophysical studies report that the contrast effect is perceived less intense for smaller patches [60–64].

**Fig 6.**
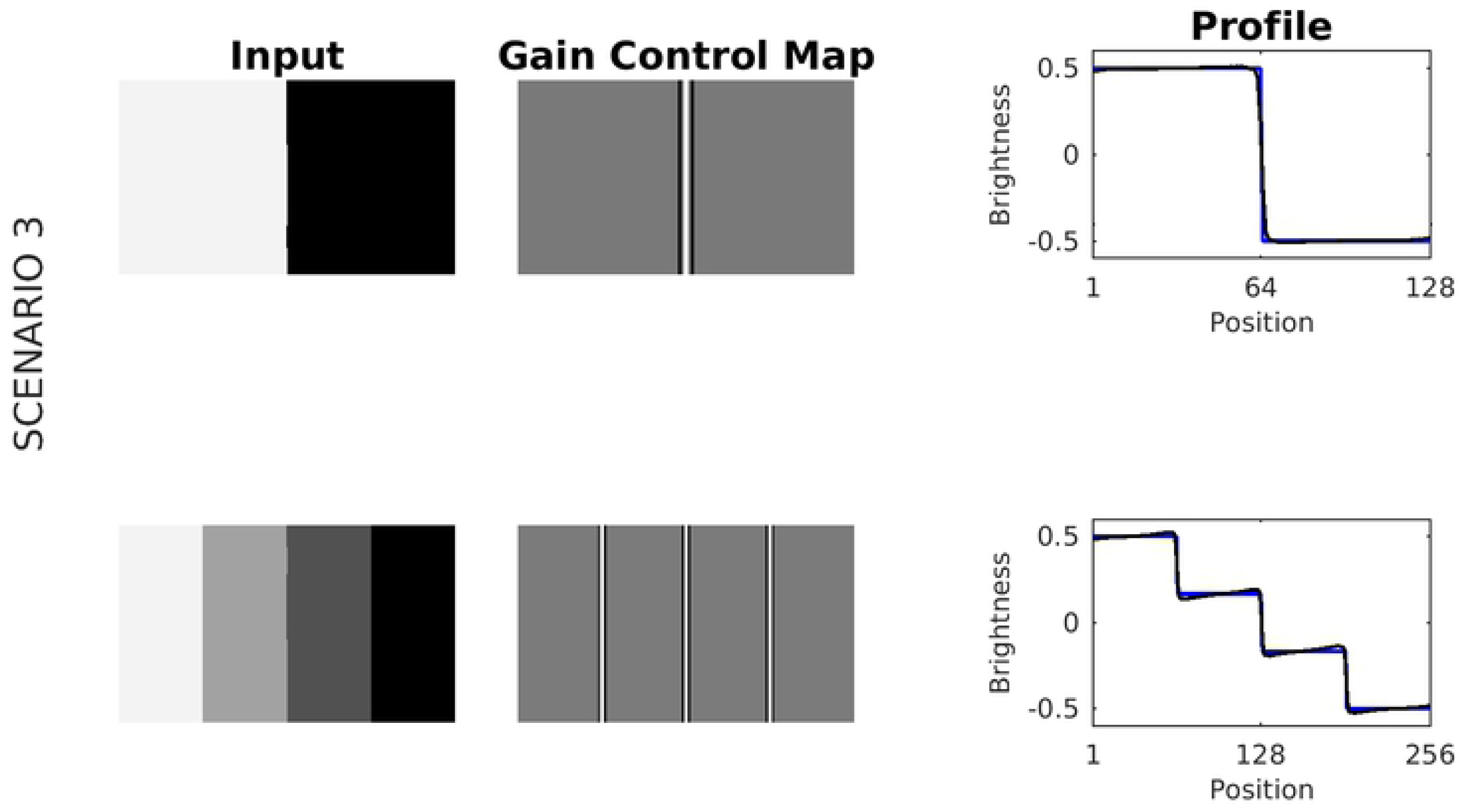
Scenario 3. A luminance step and a luminance staircase, respectively, served as input images. Activity in response to the luminance step is close to the control parameter of Eq 4 (*τ* = 0.5), producing barely changes in the corresponding brightness estimation at the edges. In contrast, for the luminance staircase, the activity at the edges is relatively far from the control parameter, inducing a boost (an increment of brightness contrast) in the corresponding brightness estimation.

**Fig 7.**
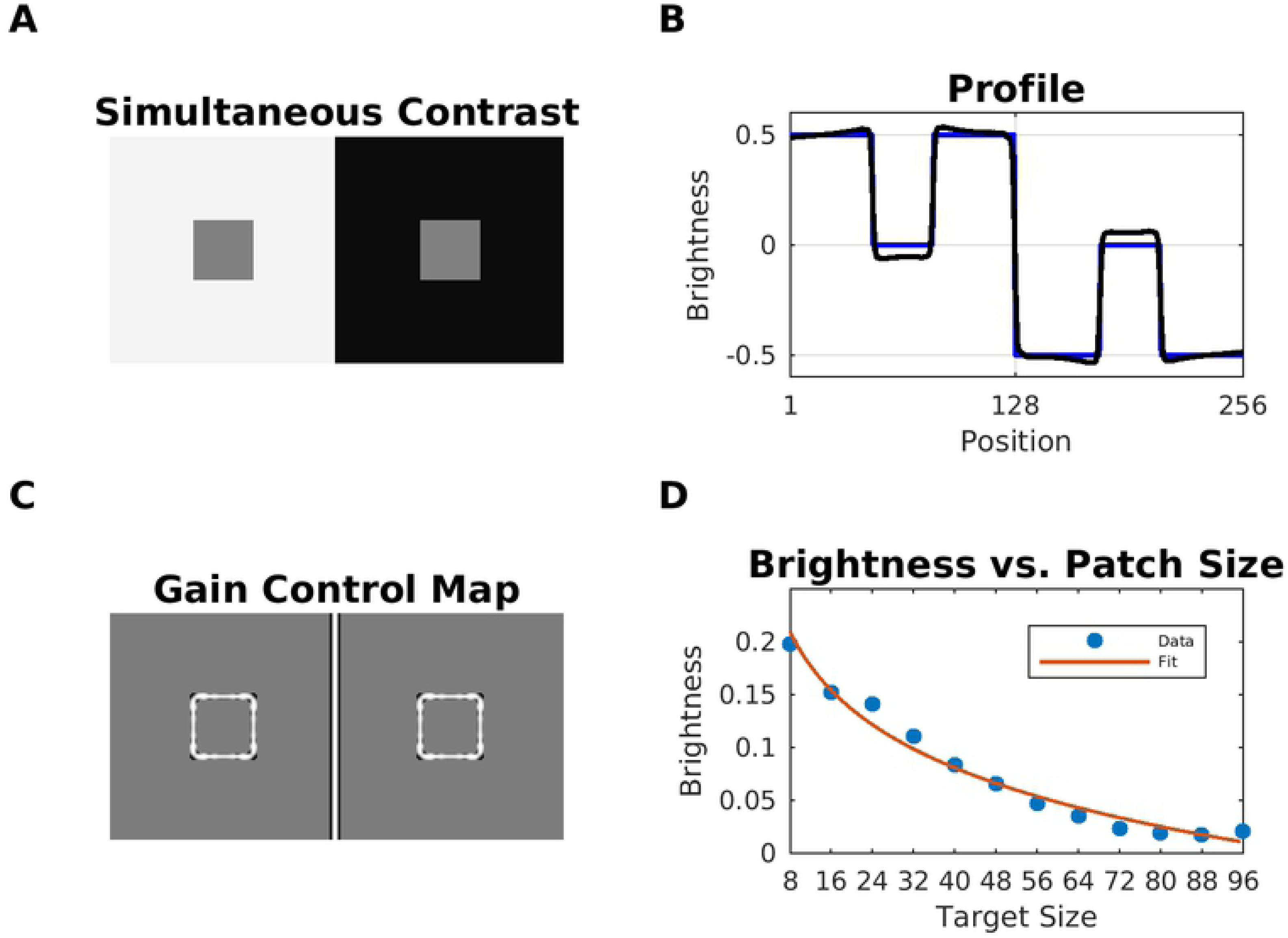
Model Prediction for Simultaneous Brightness Contrast (SBC) (A) Simultaneous brightness contrast display (model input). (B) The corresponding Gain Control Map. (C) Profile plot of the estimated brightness map (black line) and the input (blue line). (D) The quantitative relation between predicted brightness and patch size. The fit was carried out by linear regression and with the fitting function *y* = *a* + *b* log(*x*). Fitting parameters: intercept *a* = −0.3851, slope *b* = −0.0823, *R*^2^ = −0.9831, and *RMSE* = 0.0086.

Fig 7 shows the estimated brightness for SBC. In our model, the effect conforms to Scenario 1 (Fig 5). In SBC, the patterns with low spatial correlation are the patch edges with equal intensity, what causes an enhancement of their contrast after gain control (Gain Control Map in Fig 7). This translates to an increased contrast in predicted brightness (profile plot in Fig 7B). We also studied the relation between patch size and their predicted brightness. In agreement with previous studies, Fig 7D shows a logarithmic relationship between patch size and our brightness estimation [65].

#### Benary-Cross

The Benary-Cross [66] is composed of a black cross and two gray triangles with the same luminance (Fig 8). The triangle embedded in the cross is perceived as brighter than the other. Notice that both of the triangles are made up of identical contrast polarities – one white to gray and two black to gray. This effect cannot be explained by lateral inhibition and is usually attributed to “belongingness theory”, where the region in which the triangle appears to belong to induces a contrast effect [66]. Noise masking experiments support the idea that the effect is caused by low-level mechanisms [67]. Our model predicted the brightness difference in the triangles according to two scenarios. According to Scenario 1 (but also 2, considering the intensity differences), the redundant patterns correspond to those edges of the triangles which are aligned with the cross. They latter are attenuated, while the non-redundant edges are enhanced. The Gain Control Map suggests a rather balanced effect, what is confirmed by the model’s predicted brightness map. Notice, however, that the length of the non-suppressed edges is bigger for the left triangle.

**Fig 8.**
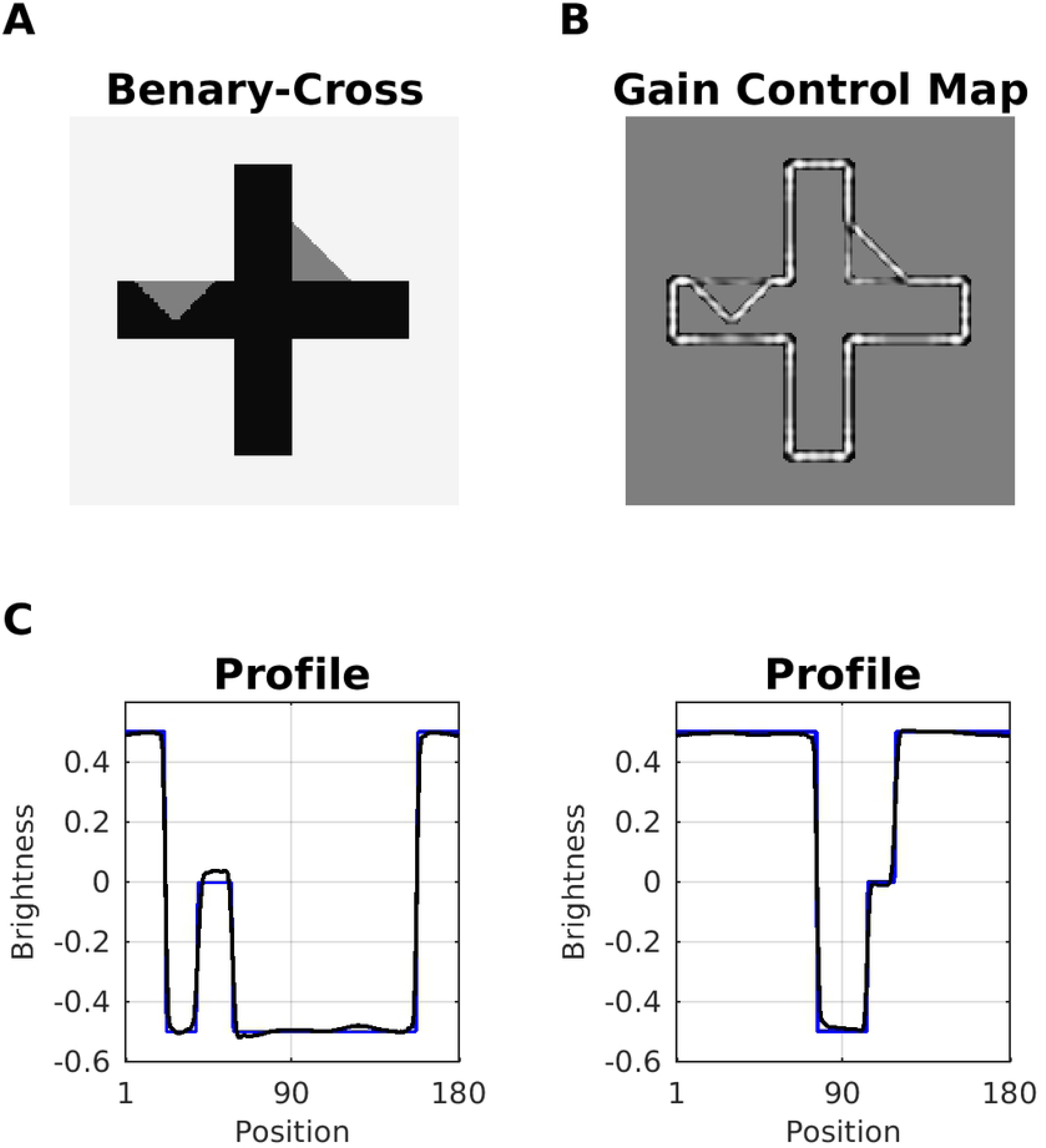
Model Prediction for Benary-Cross illusion. (A) Benary-Cross. Both triangles have the same intensity, but the triangle embedded in the cross is perceived as brighter. (B) In the gain control map, redundant edges (=aligned with the cross) of the triangle are weakened. (C) Profile plots of predicted brightness (blue line) versus luminance (black). The left profile plot shows the left triangle.

#### Reverse Contrast

Reverse Contrast (Fig 9) was introduced by Gilchrist in the context of his anchoring theory. Gilchrist and co-authors suggest that simultaneous brightness contrast (SBC) can be reversed (e.g. by overcoming lateral inhibition) by adding more structures to the original SBC display. The purported mechanism acts on grounds of perceptual grouping of these structures.

**Fig 9.**
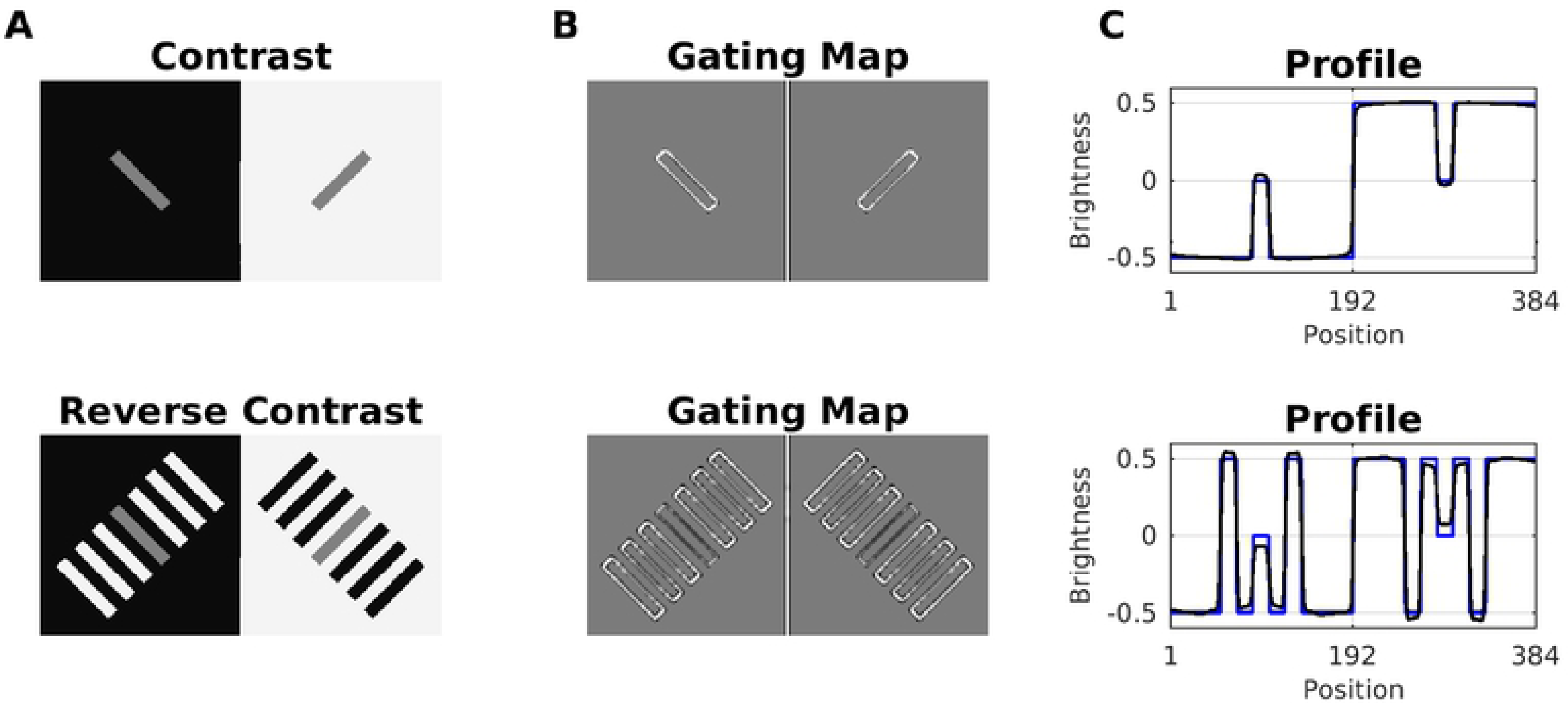
Model Prediction for reverse contrast effect. (A) Simultaneous Brightness Contrast (SBC, top) and Reverse Contrast (bottom) which is constructed by adding flanking bars to the SBC configuration. All gray patches have the same luminance. Reverse contrast can be explained either by assimilation with the in-between bars that have the same intensity as the background, or as contrast with the flanking bars that have the opposite luminance to the background. (B) Gain Control Maps obtained by dynamic filtering. Notice the suppression of parallel edges corresponding to flanking patches with the same intensity. (C) Profile plots of predicted brightness (black line) versus luminance (blue).

Our model predicted the reverse contrast effect according to Scenario 1 and 2, respectively. In case of SBC, dynamic filtering increments the activity of the non-redundant edges, which outline the two patches (Gain Control Map in Fig 9). In the case of reverse contrast, the redundant activity depends on both edge orientation (Scenario 1) and contrast polarity (Scenario 2). Accordingly, all parallel edges of the (flanking) patches with equal intensity are weakened by the dynamic filter. However, the central patch has a different intensity than the flanking patches, and its edges are enhanced.

In order to better understand how our model predicted reverse contrast, we probed it with further configurations (see Fig 9). We observed that the change in brightness of the gray patches increases as a function of the number of flanking bars (Fig 10A). On the other hand, if the flanking bars were misaligned to various degrees (disrupting the good continuation principle of perceptual organization), the effect was considerably reduced (Fig 10B). Both results stand in agreement with psychophysical experiments [68]. However, in the latter study the authors examined displays with even more configurations that our model cannot predict (results not shown).

**Fig 10.**
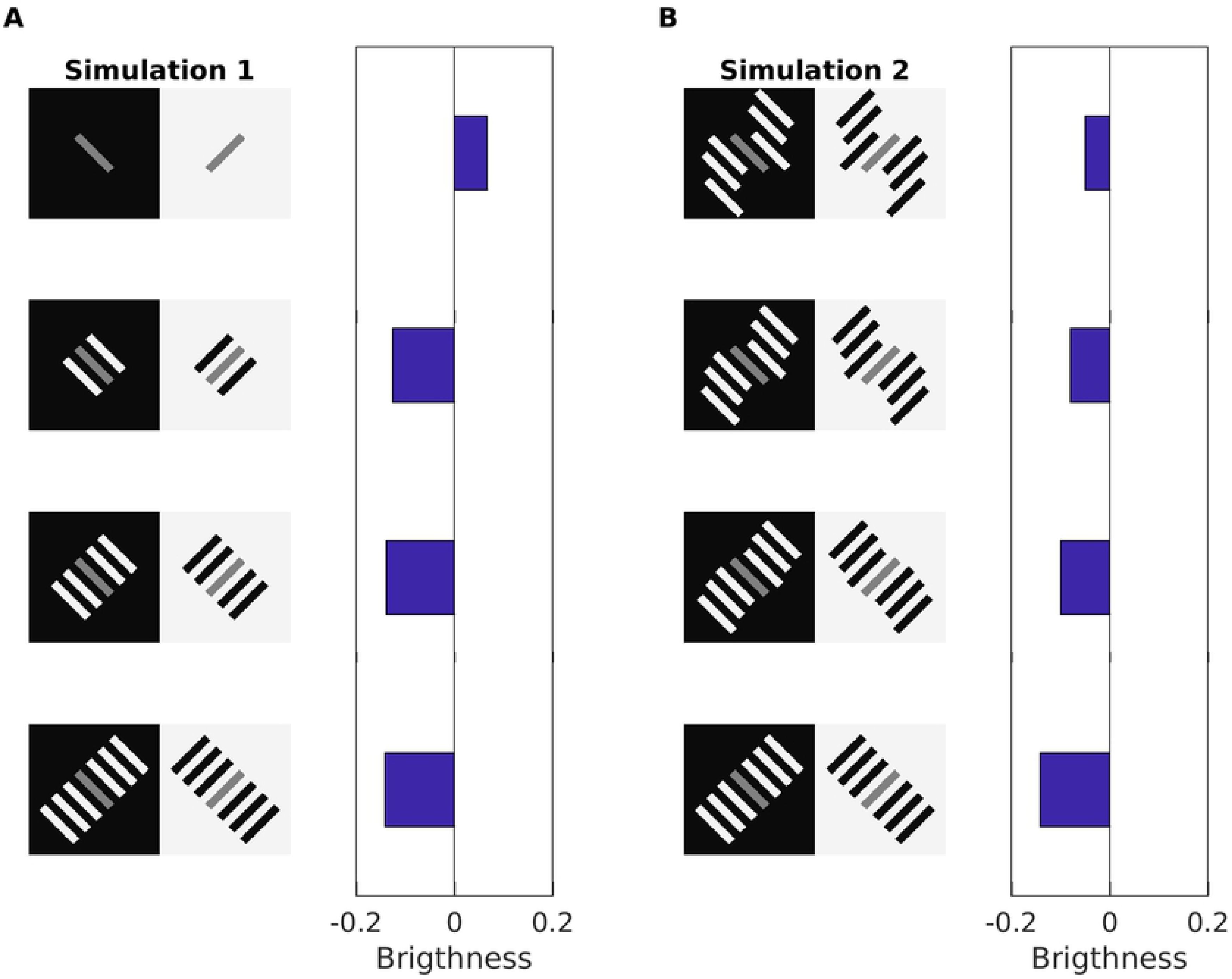
Model Prediction for reverse contrast effect for displays with different configurations. (A) Reverse contrast with a varying number of adjacent bars to the gray patch. The bar plots show the predicted brightness difference between the gray patches for the corresponding display (a positive value indicates contrast, while negative values “reverse contrast”). (B) Reverse contrast where the good continuation of the end points is varied. This in turn affect the suppression of redundant edges, which increases with the alignment of the flanking bars.

### Brightness Assimilation Effects

#### White’s effect

Fig 11A shows the White’s effect, where two gray bars with identical luminance are embedded in alternating black and white stripes. The bar on the black stripe is perceived as brighter as the other one. Lateral inhibition cannot account for this effect, and it has been suggested that the effect is caused by assimilation [3, 69, 70]. Assimilation means that the brightness of the flanking stripes averages with the gray bars, and therefore one expects that reducing the bar height would also reduce the strength of assimilation. However, experimental data indicate that the perceived difference between the bars increases with smaller heights [71], and that bandpass-filtered noise with the same orientation as the stripes enhanced the effect, while with perpendicular orientation the effect was diminished [67]. Therefore, White’s effect seems to be principally generated by contrast at the horizontal edges of the bars (Fig 11B and Fig 11C), and to a less detect by assimilation from the flanking stripes [72]. In fact, a mainly contrast-based account is supported by the Gain Control Maps of Fig 11A and Fig 11B. Because the vertical edges (assimilation) are highly redundant, their activity is diminished (Scenario 1 & 2). The brightness estimation is dominated by the horizontal edges of the bars (contrast), which are enhanced. The vertical edges nevertheless account for a residual assimilation, but the effect in estimated brightness is less for Fig 11B than for 11A. Therefore, the display with the smaller bars (Fig 11B) has a higher predicted brightness difference between them, because less activity from the vertical edges “mixes” with that from the horizontal edges during filling-in.

**Fig 11.**
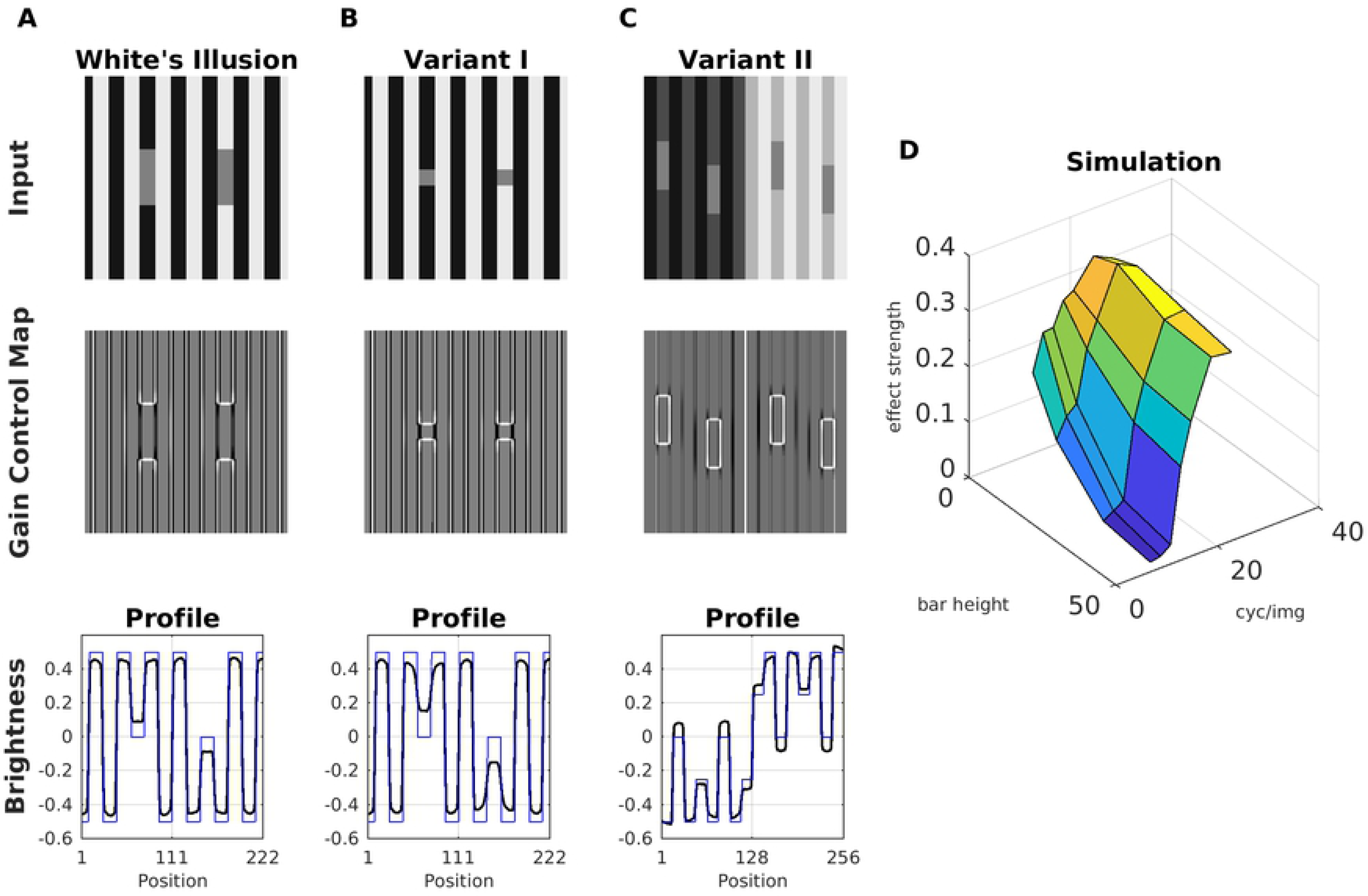
Model Prediction for White’s Illusion. (A) Top: White’s illusion; middle: the corresponding gain control map; bottom: profile plot of estimated brightness (black line) and luminance (blue line). (B) With smaller bar height, the brightness difference between the bars increases. (C) Modification of White’s illusion which produces a strong contrast effect. (D) Surface plot of the estimated brightness difference (effect strength) between the bars as a function of bar height (in units of pixels) and spatial frequency of the background stripes (in units of cycles/image). Image size was 256 x 256 pixels.

Despite of the presence of stripes (but with low contrast), the display of Fig 11C shows a clear contrast effect of the bars (cf. Fig 11C) according to Scenario 2. We also studied the relation between the target size and the brightness estimation. We observed that the predicted brightness of the bars could be modified as a function of bar height and spatial frequency of the background (Fig 11D). Specifically, the predicted brightness difference between the bars increases both with decreasing bar height and with increasing spatial frequency. These model predictions are thus in agreement with previous studies [10].

#### Todorovic’s illusion

Todorovic’s illusion consists of a display with two luminance disks with identical intensity and two sets (black and white) of four squares. The original illusion is designated as Context B in Fig 12, where the disks are occluded by the squares. The test patch occluded by the white squares appears to be brighter than the other. This illusion was originally explained in terms of T-junctions [26]. However, [23] showed that the effect persisted without T-junctions. Later, [73] studied different variations of Todorovic’s Illusion (labeled Context A and Context C in Fig 12). They found that the target size interacted with the strength of assimilation. The original effect can be reversed according to Context C (Fig 12), which looks like looking through a window cross. It can also be abolished by moving the disk into the foreground (Fig 12, Context A). The bottom row of Fig 12 shows the profile plots of the brightness maps produced by our model. The results could be understood by analyzing the corresponding gain control maps. For Context B, the activity of the occluding edges (disk with squares) was reduced by dynamic filtering. As a consequence, less contrast is produced in the brightness estimation. On the other hand, the edges between the disk and the (black or white) background was enhanced, what produces more contrast in the brightness estimation. This double effect combined to the finally predicted brightness. For Context C, the effect was analogous to Context B, but with opposite disk brightness. With Context A, edge activity along the disks’ circumferences was enhanced. Note that the edge covers the squares as well as the background. Although this enhancement of activity produced locally more contrast, it is approximately the same for the two disks, thus producing almost no effect (profile plot of Context B). Fig 13 shows simulation results for the three Contexts as a function of disk size, where we observed qualitatively similar results to previous studies [73].

**Fig 12.**
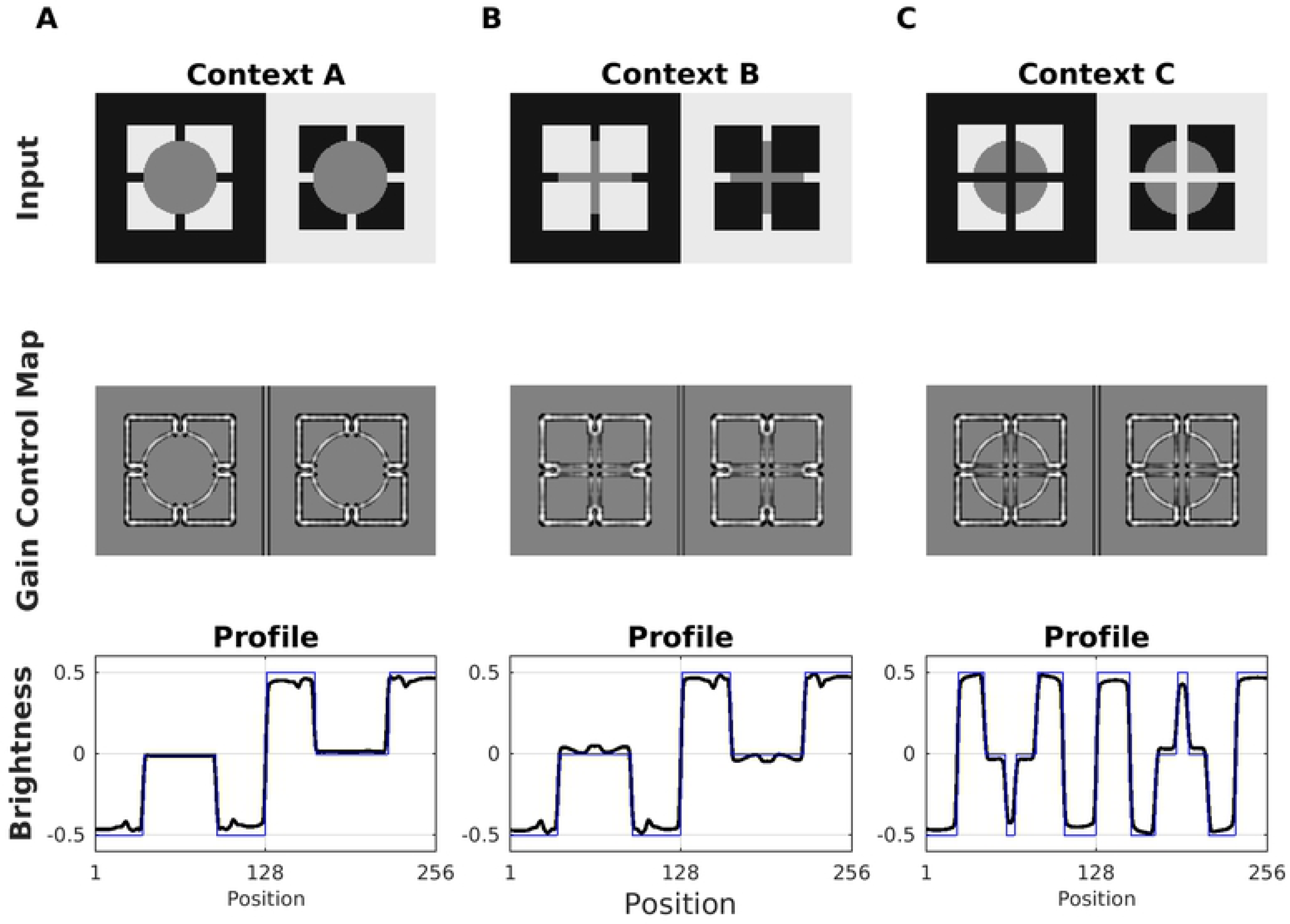
Model Prediction for Todorovic’s illusion. (A) Top: A variation of Todorovic’s original illusion with the gray disks in the foreground (Context A). An effect is hardly perceivable. Middle: The corresponding Gain Control Map. Bottom: Profile plots of estimated brightness (black line) and luminance (blue line). The model predicted at most a very weak effect. (B) Original Todorovic Illusion (Context B), where the occluded left disk is perceived as being brighter. (C) Reversed Todorovic Illusion (Context C). Now it looks like viewing the disks on a single square background through a window cross, and the left square is perceived as being darker.

**Fig 13.**
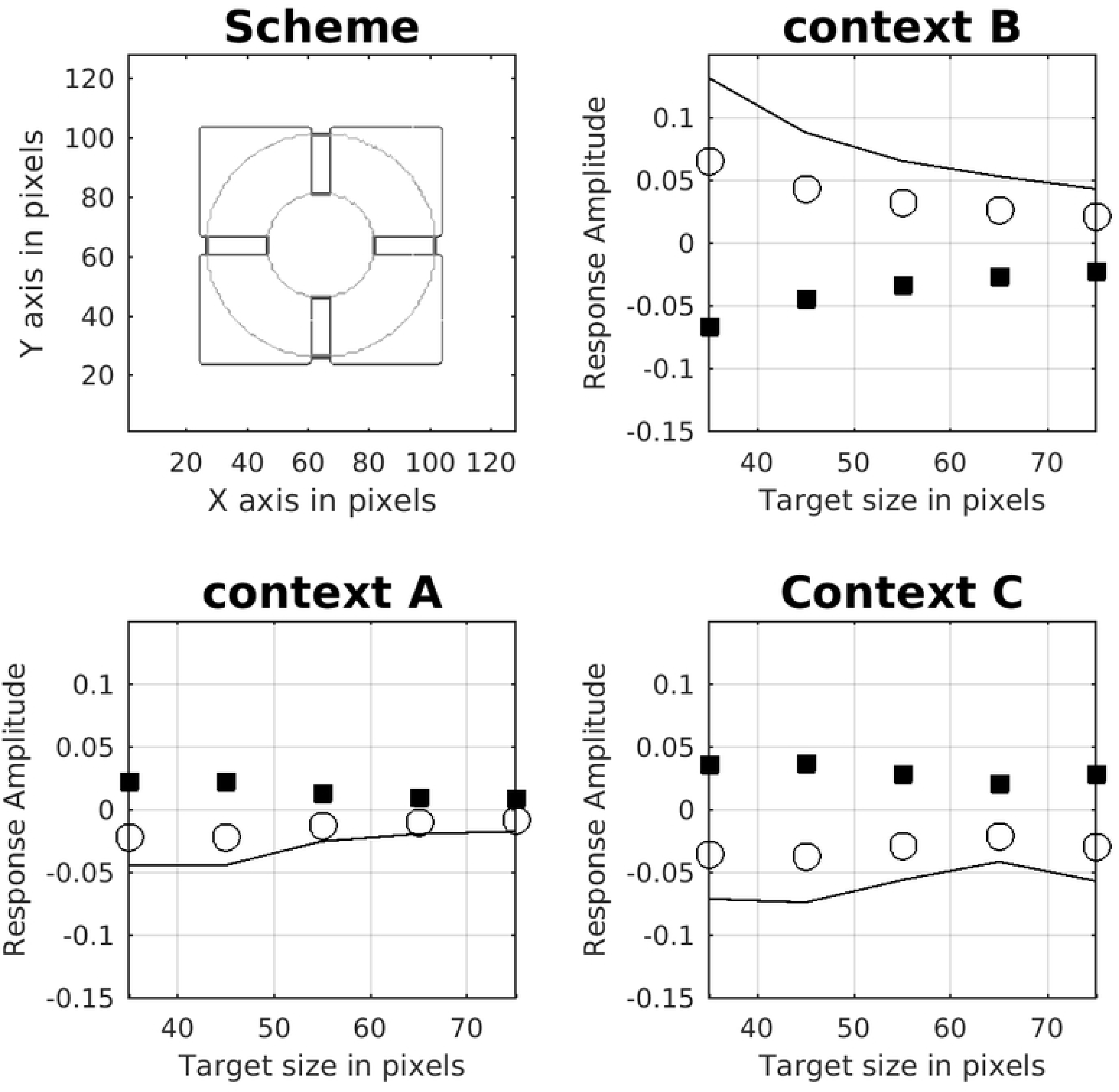
Brightness dependence on target size for Todorovic’s display. The Scheme shows the smallest and biggest disk size that was used with respect to the squares in order to generate the plots. Each plot indicates the brightness effect for each of the three Contexts shown in Fig 12. The empty circles indicate the predicted brightness of the disk with the white squares. The filled symbols show disk brightness with the black squares. The continuous lines show the estimated brightness difference between both disks and indicate the predicted strength of the illusion.

#### More assimilation displays: Dungeon, Checkerboard and Shevell

The predictions of our model generalize well to further assimilation displays. Figure 14 shows the Dungeon illusion [25], the Checkerboard illusion [8] and Shevell’s Ring [74]. Although all three reveal an assimilation effect on the gray areas, they are different with respect to their spatial configuration. Notice in particular the absence of T-junctions in the Shevell’s Ring display. Fig 14 also shows the corresponding simulation results. All three illusions can be explained according to Scenario 2 since the edges corresponding to the gray areas represent redundant patterns with low intensity.

**Fig 14.**
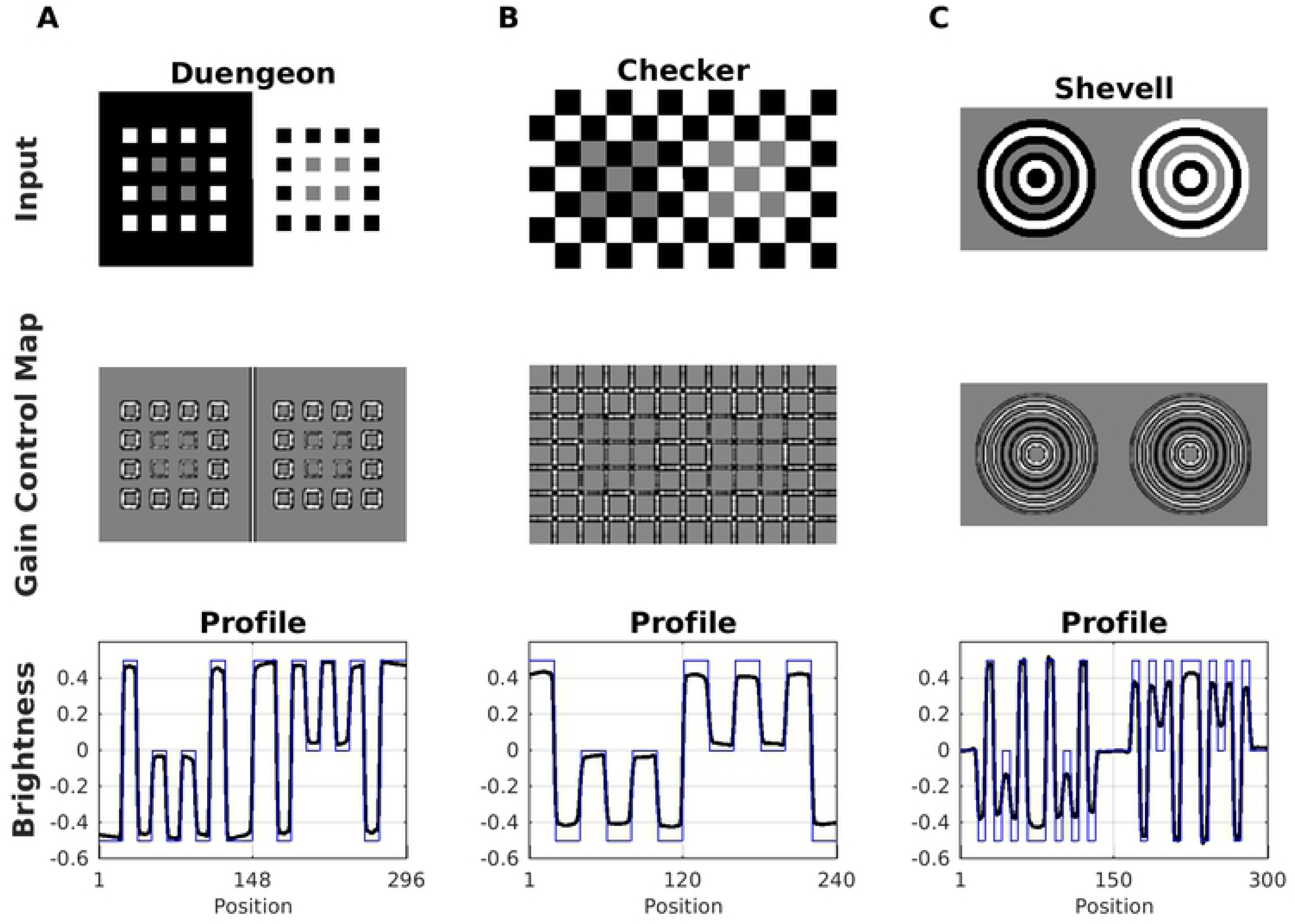
Model Predictions for Dungeon, Checkerboard and Shevell. In each display, the gray areas have the same luminance, yet they are perceived differently because of assimilation with the adjacent structures. (A) Top: Dungeon illusion. Middle, corresponding gain control map. Bottom, profile plot of the estimated brightness (black line) compared with the input (blue line). (B) Checkerboard illusion. (C) Shevell’s Rings. Notice that this illusion cannot be explained with T-junctions.

### Further Visual Illusions

#### Craik-O’BrienCornsweet Effect (COCE)

Fig 15A shows the COCE, which consists of regions separated by opposing luminance gradients (“cusps”) starting at the edges. The cusps drop quickly to a homogeneous gray level and thus the regions between the edges have the same luminance. Nevertheless, especially at low contrast, the gradients seem to fill into the intermediate regions, such that the display is perceived as a low-contrast rectangular wave. The perception of a rectangular wave is less pronounced with high contrast cusps.

**Fig 15.**
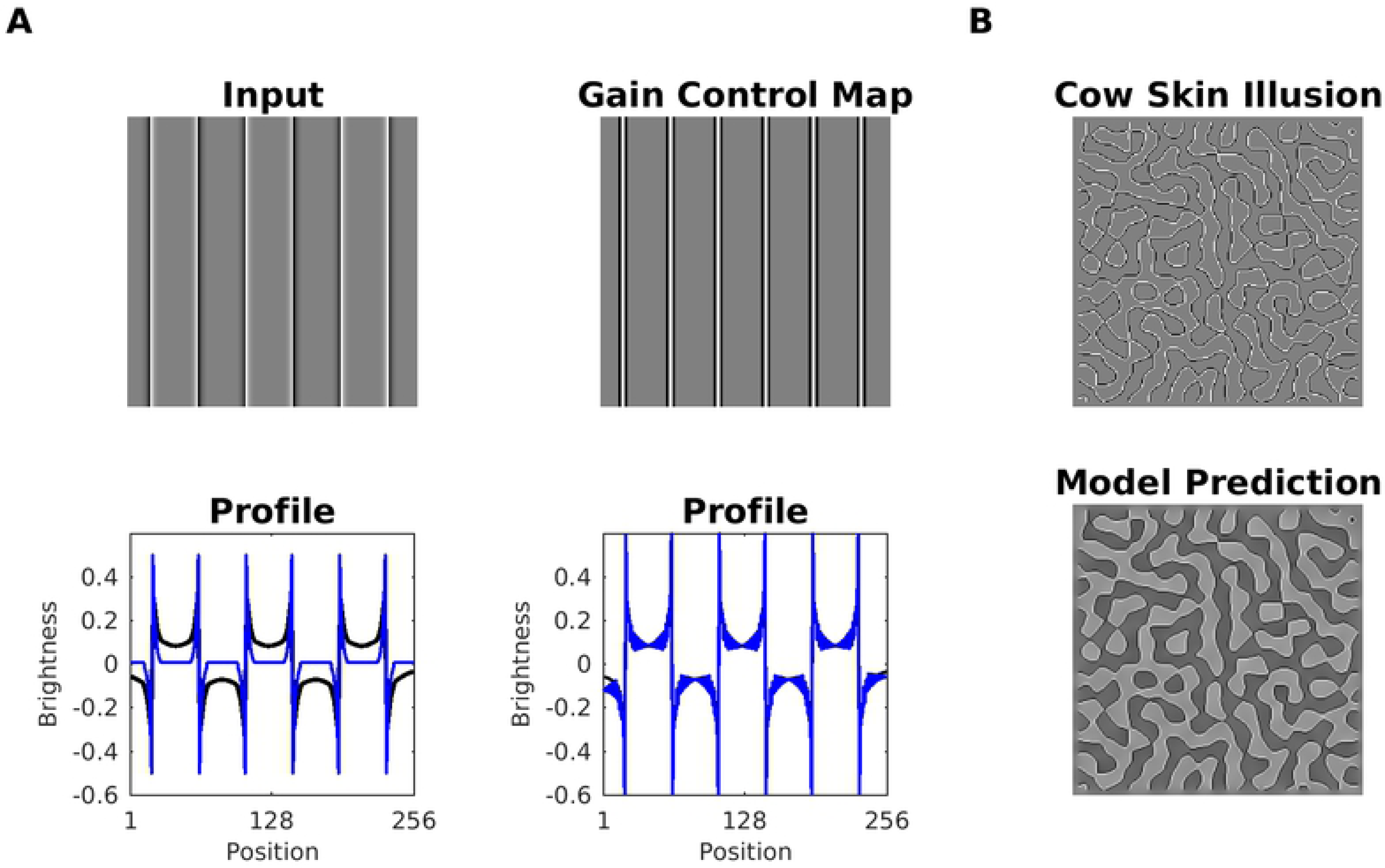
Model Prediction for Craik-O’BrienCornsweet Effect (COCE) (A) Top: COCE along with the Gain Control Map. Notice that black lines (adjacent to the edges) in the Gain Control Map, which indicate negative values. Bottom: The first profile plot shows the predicted brightness (black line) along with input luminance (blue line). The second profile plot shows the predicted brightness without low-pass filtering after the gain control mechanism (Eq4). (B) The cow-skin illusion is a variant of the COCE without luminance gradients. It consists only of adjacent black and white lines. Our model consistently predicted this illusion: The brightness map generated by our model is shown at the bottom right.

Our model predicted the COCE, but the explanation is more intricate. At first sight, being a filling-in type effect, it should be predicted in a straightforward way by our model. We verified that without gain control (Eq4), the COCE cannot be predicted (data not shown). Could it be then that the effect is produced by the low-pass filtering which is applied after the gain control mechanism (see method section). This is not the case, since the removal of low-pass filtering did not affect the prediction of the COCE (see profile plot 2 in Fig 15A). Therefore, the gain control mechanism contributes to producing the effect. Indeed, the luminance gradients cause negative values (indicated by black lines) in the Gain Control Map around the edges (cf. Fig 15A). As a consequence, activity corresponding to the luminance gradients is suppressed by the gain control mechanism, what furthermore reduces the peak activity at the edges. After all, the gradients are “ignored”, and our model generates the COCE as a result of assimilation of the edges. This explanation is also consistent with the cow-skin illusion, which is a variant of the COCE without luminance gradients. It is composed exclusively of adjacent black and white lines, and the empty regions are randomly arranged. Fig 15B shows the brightness prediction for the cow-skin illusion.

#### Hermann/Hering grid

Fig 16A (in the top) shows the Hermann/Hering Grid (HG). Although the luminance between the black squares is constant, illusory gray dots appear at the (white) intersections. The textbook explanation deems the center-surround receptive fields of retinal ganglion cells as the principal mechanism [75]: Assume a circular receptive field with an excitatory center which has the same width as the white grid lines. The inhibitory surround covers in addition the black squares. If the center is located right at an intersection, it receives more inhibition from the surround (from two white lines) than when the center is positioned between two intersections (inhibition from one line). This translates to a brightness reduction at the intersections, but not in between. This mechanism, though, is insufficient to explain why the effect is considerably reduced (or even removed) if the bars are slightly corrugated (Fig 16A, bottom). It is also reduced as a function of the ratio between grid line width and block width, where no effect is produced for a ratio of one [76].

**Fig 16.**
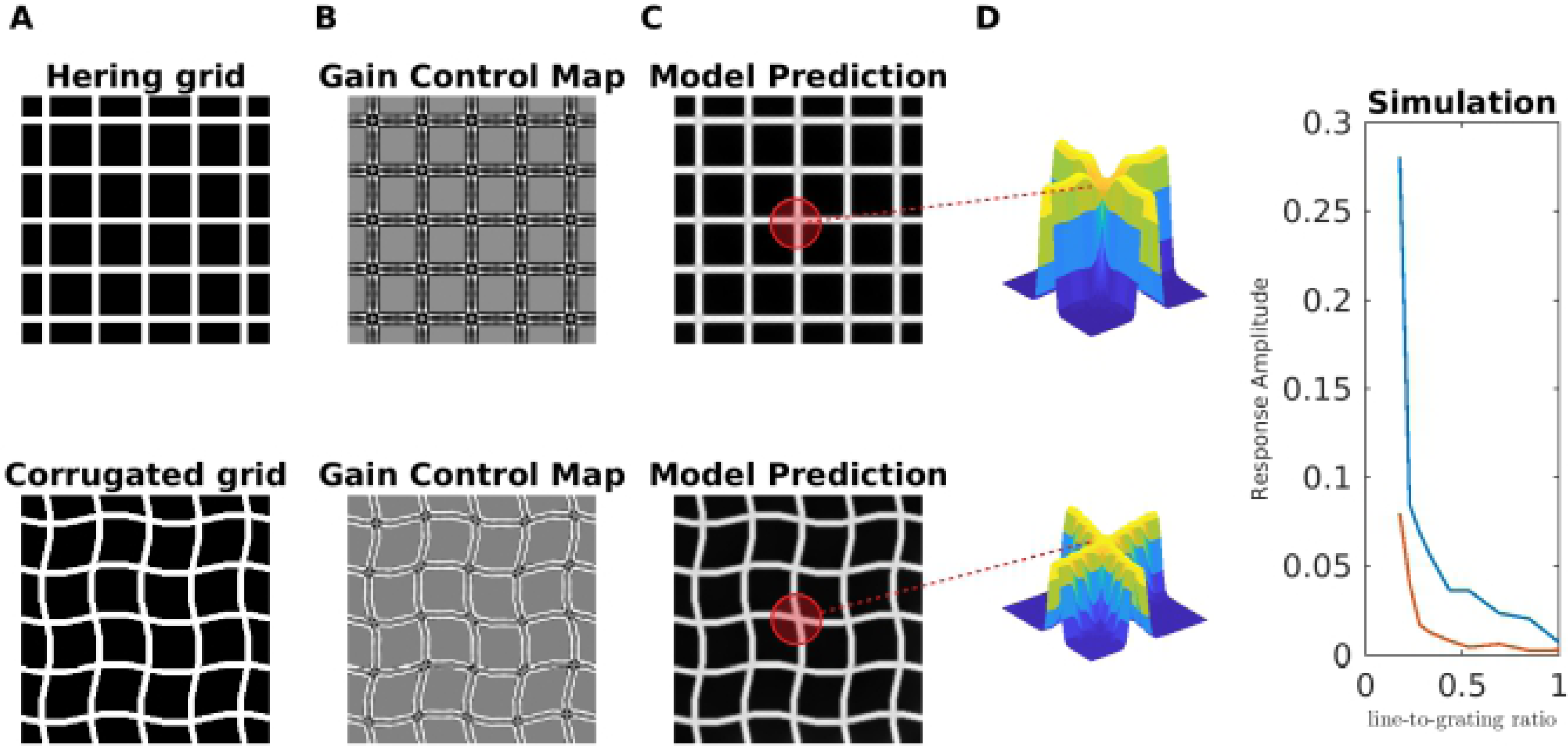
Model Predictions for Hermann/Hering grid and corrugated grid. (A) Hermann-Hering (HG) illusion and a corrugated version of it. At the intersections of the white grid lines, illusory gray spots are perceived in the HG, but not in the corrugated grid. (B) The corresponding gain control maps of the input images of A. (C) The brightness estimation from our model. The surfaces plots (insets) illustrate the 3D profile of the brightness estimation corresponding to regions highlighted with red. (D) The predicted brightness magnitude at the intersections as a function of the ratio 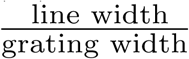, where the red curve corresponds to the corrugated grid, and the blue curve to the HG.

Our model predicted the darkening of the intersections for the HG, and the absence of darkening for the corrugated HG. The HG adheres to Scenario 1, where redundant activity is inhibited. Inhibition is especially strong at the intersections of the Gain Control Map. In this way, an assimilation effect is induced. As to the corrugated HG, the corners also represent the redundant patterns, but because the spatial structure are less regular, the inhibition is correspondingly weaker (compare the Gain Control Maps shown in Fig 16B). Consequently, the brightness reduction at the intersections is considerably weaker for the corrugated HG.

Fig16D shows the dependence of the darkening effect on the ratio between grid line width and block width. In agreement with the results from [76], we find that the darkening effect decreases while the ratio approaches one. We also wish to note that we were unable to predict further results with the HG that were presented in [76].

#### Luminance Staircase and Pyramid (Chevreul’s illusion)

Chevreul’s illusion consists of increasing levels of luminance, arranged as a staircase or as a pyramid. Although luminance is constant at each step, one perceives an illusory brightening on the side of each step where the adjacent step is darker, and an illusory darkening on the other. In the pyramid version, one perceives in addition (illusory) glowing diagonals (Fig 17). The effect is absent at the lowest (black) and highest (white) luminance level, and is considerably reduced on the middle step for a staircase made up of three steps.

**Fig 17.**
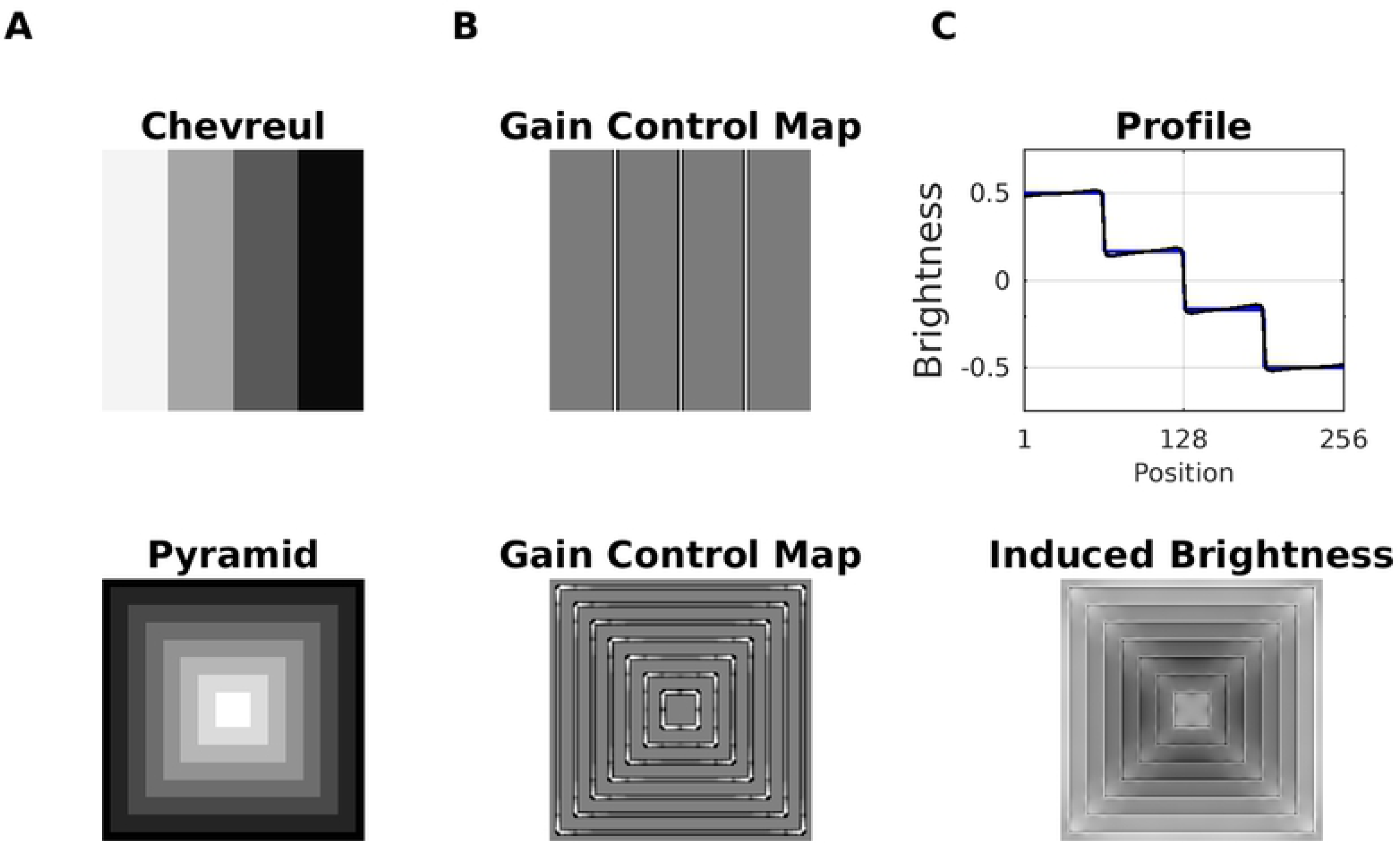
Model Predictions for Luminance Staircase and Pyramid (Chevreul’s illusion) (A) Top: Luminance staircase. Bottom: luminance Pyramid. (B) The corresponding gain control maps as a result of dynamic filtering. (C) Top: Profile plot of the estimated brightness (black line) of the luminance staircase (blue line). Bottom: The induced brightness, which consists of the difference map between estimated brightness and input luminance (i.e., brighter gray level mean positive values, and darker gray levels mean negative values).

All aspects of Chevreul’s illusion are consistently predicted by the gradient system, which is a computational model for representing luminance gradients [77, 78]. The idea behind gradient representations is to capture the smooth variations of luminance (illumination effects), in order to help to disentangle reflectance from the illumination component in luminance (since luminance is the product of reflectance with illumination).

Brightness predictions from our model for the luminance staircase and the pyramid are shown in Fig17. The illusory whitening and darkening at the stairs can be explained according to Scenario 3: On the one hand, the gain control map increases the activity of the Contrast-Luminance channel. On the other hand, the increase in excitation is offset by the normalization mechanism, thereby producing non-uniform brightness activity at the stairs. The glowing diagonals of the Pyramid Illusion are produced according to Scenario 1, where the activity of non-redundant spatial patterns – especially at the corners – is enhanced. On the other hand, the edges of the staircase represent a redundant pattern, the activity of which is decreased. Consequently, more (less) contrast at the corners (at edges) is generated in the brightness estimation (Fig 17C, bottom). Finally, it is essential to highlight a limitation of our model in this context. We observed that for a big number of steps (i.e., very narrow steps) the dynamic filter “collapses” and the model could not longer predict the illusion nor the glowing diagonals (data not shown). This is a consequence of the scale-sensitivity of the dynamic filter (i.e., the size of the sampling patches), since with decreasing step size, the staircase eventually approaches a linear luminance gradient, and the filter cannot resolve anymore individual steps.

#### Mach Bands

Mach Bands are illusory glowing stripes that are perceived adjacent to knee points that are connected with a luminance ramp, where the bright (dark) band is attached to the plateau with high (low) luminance (Fig 18). Notice that Mach bands do not cause Chevreul’s illusion. The perceived strength of Mach bands decreases when the ramp gets steeper and eventually approaches a luminance step. Also, for very shallow ramps, the perceived strength decreases. The perceived strength has thus a maximum at intermediate ramp widths [79, 80]. The textbook explanation based on lateral inhibition is insufficient to explain the variation of strength with ramp width – it would wrongly predict maximum perceived strength at a luminance step [81, 83, 84]. The perceived strength of Mach bands is also modulated by the proximity, contrast and sharpness of an adjacently placed stimuli [81, 84].

**Fig 18.**
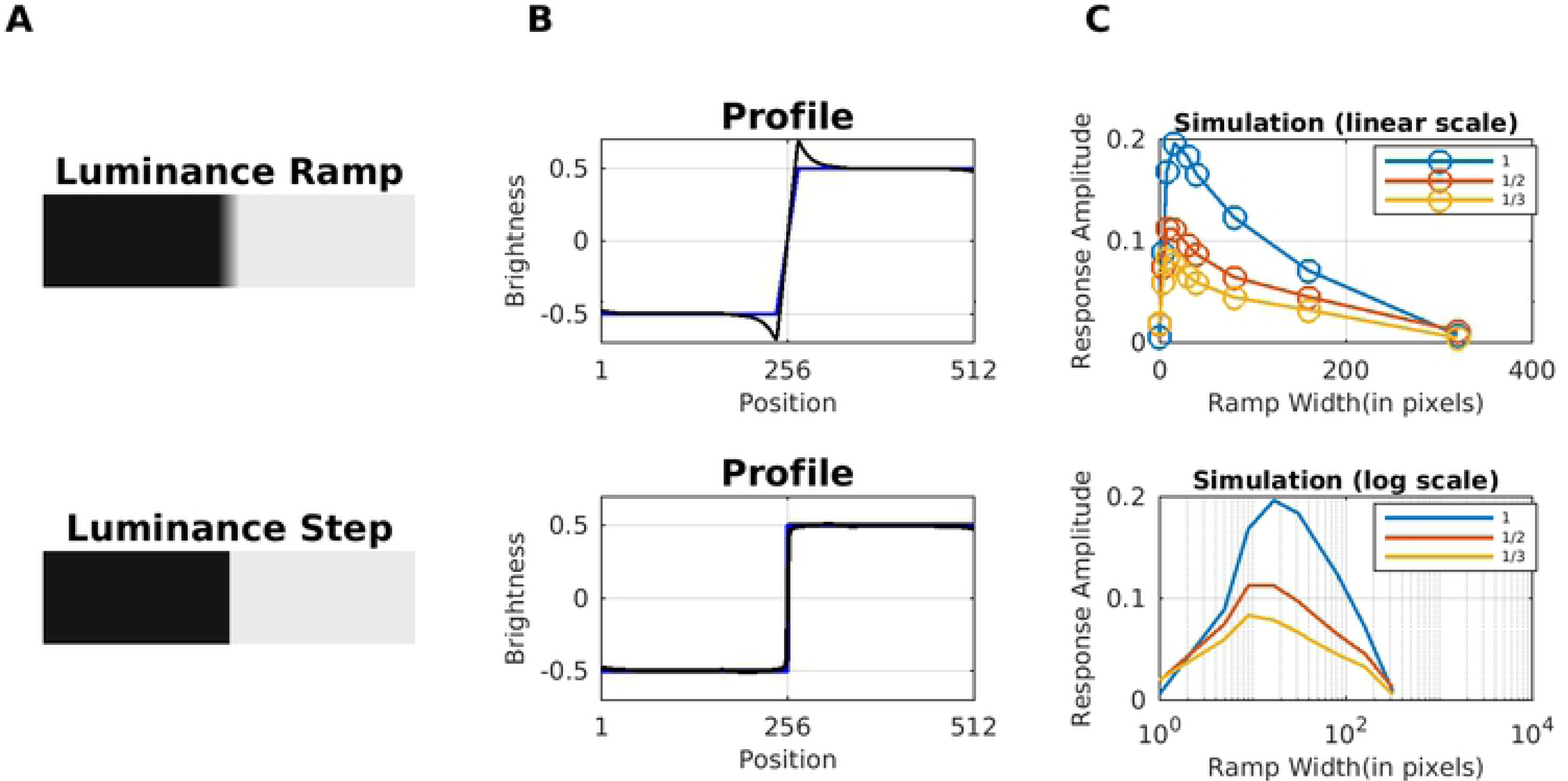
Model Prediction for Mach Bands. (A) Top, A luminance ramp that leads to the perception of Mach bands close to the knee points of the ramp. Bottom, a luminance step (no Mach bands are perceived). (B) Profiles plots of estimated brightness (black line) compared with the corresponding input (blue line) of A. (C) Brightness magnitude (at the knee point of the ramp) as a function of ramp width. The plots show the predictions of our model on the perceived strength of the bright Mach band. In the top plot, the abscissa is scaled linearly, while it is scaled logarithmically in the bottom plot. The different curves represent different dynamic ranges (i.e., differences between luminance values of the upper and the lower plateau).

The only computational model published so far that quantitatively predicted all published data about Mach Bands is the gradient system [77]. The gradient system suggests that Mach bands are also perceived at the peaks and troughs of a triangular wave. The gradient system furthermore suggests that bright Mach bands are key for the perception of light-emitting surfaces [78]. Our model predicted the Mach bands, and as well as the absence of them at steps (see profile plots in Fig 18). It furthermore succeeds in predicting the inverted-U curve of the perceived strength of Mach Bands as a function of the ramp width (Fig 18C). The inverted-U curve could be explained by two mechanisms which act in opposite ways. (i) If the ramp width decreases, then the activity at the knee points reaches a maximum that renders the normalization mechanism (denominator of Eq 4) of the gain control mechanism ineffective (Scenario 3; ideal step luminance). If the ramp width increases, then the luminance transition between the plateaus is more gradual, which is associated with less activity at the edge locations. In this way, the edge activity gets more susceptible to gain control mechanism towards the maximum of the perceived strength. However, this effect does not remain constant. After a certain ramp width, the activity across the ramp gets comparable to the activity at the knee points, which produces less variability in the energy map E. In consequence, (ii) the dynamic filter has less effect in eq 2, reducing gradually the perceived strength induced by the dynamic filtering.

#### Grating Induction (GI)

Fig 19 shows the grating induction (GI) display [85], which consists of two sinusoidal gratings (inducers) separated by a gap (test field). Although it has a constant luminance, an illusory brightness modulation is perceived across the test field if the two inducers are in-phase. The brightness modulation has the opposite phase as the inducers, and the effect decreases when shifting the phase of the inducer gratings relative to each other, being minimum when the gratings are in anti-phase. The illusory modulation is furthermore attenuated with increasing distance between the inducer gratings and with increasing spatial frequency. The GI can be explained in terms of multi-scale filtering [9, 60]), but also by filling-in models [86]. Notice that a common misconception with diffusion-based approaches (=filling-in models) is that the illusory brightness modulation across the test field would average out. This, however, is usually not the case. The exact explanation depends on the model under consideration. For instance, a mechanisms that counteracts “averaging out” are boundary webs from the boundary contour system (BCS) that extend across the test field and trap feature contour activity (FCS) [87]. Other mechanisms include cross-channel inhibition between brightness and darkness activity during filling-in [88].

**Fig 19.**
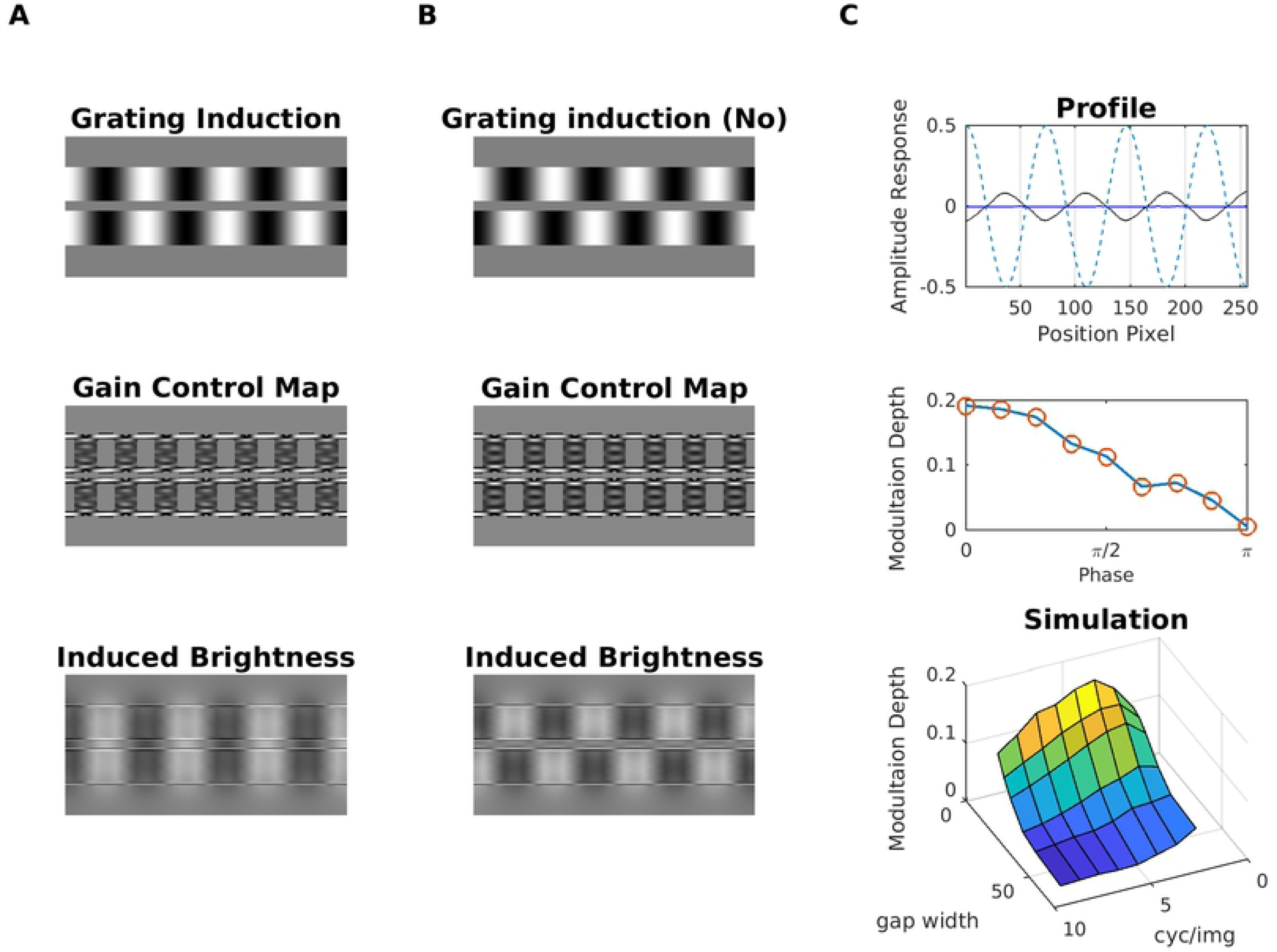
Model Prediction for Grating Induction. (A) The Garing Induction refers to the illusory perception of a brightness modulation across the gap (=test field) between the inducer gratings. The brightness modulation is perceived in opposite phase to the inducer gratings. (B) When the inducer gratings stand in opposite phase to each other, then brightness modulation is considerably reduced. The corresponding gain control maps are shown in the middle row, and the last row shows the induced brightness, which consists of the difference map between estimated brightness and input luminance (i.e., brighter gray levels mean positive values, and darker gray levels mean negative values). (C) Top: Profile plot of brightness estimation for display A (black line) and display B (blue line). The dashed blue line shows the luminance profile of the inducer grating A. Middle, modulation depth as a function of the phase difference between the two inducer gratings. Bottom: Surface plot that shows how modulation depth depends on test field width and spatial frequency of the inducer grating.

Fig 19 shows that our model consistently predicted the illusory modulation of brightness across the test field. The strength of the effect decreases in an approximately linear fashion with increasing phase difference between the inducer gratings (Figure 19C). We also observed from a specific spatial frequency on (4 cycles/image) that the brightness modulation decreases with increasing separation and spatial frequency, respectively, of the inducer gratings (surface plot in Fig 19C). Unlike the rest of the illusions the GI effect was produced mainly at the filling-in stage (Eq 5), and to a lesser degree by dynamic filtering. Dynamic filtering increased the activity at the boundaries of the inducer gratings (see gain control maps in Fig 19); this increment produces a significant contrast in estimated brightness between the inducer gratings and the test field, which eventually propagated (by filling-in) across the test field.

### Real World Images and Noise

Although synthetic images are a valuable tool for the study of certain aspects of the visual system, it nevertheless evolved to the processing of real-world images. Real-world images provide, therefore, a test of robustness for any model of the visual system. Fig 20 demonstrates that our model is capable of real-world image processing, as well as its ability to handle noisy versions of visual illusion displays (Benary-Cross, Simultaneous Brightness Contrast, and White’s Illusion). Previously we showed that by using a set derivative filters that cover all orientations (similar to simple cells), Eq 5 globally converges to a stable solution [89]. The convergence is robust against adding noise to the input, or using a high dynamic range of luminance values (Fig 20A, top, middle). The robustness extends to the consistent prediction of visual illusions in the presence of additive noise (Fig 20B). In particular, we noted that simultaneous brightness contrast was more sensitive to the presence of uncorrelated noise than assimilation displays. The robustness against noise relates to dynamic filtering, which reduces the correlated (or redundant) spatial information in the edge map. The redundancy of the edges would not be affected by spatially uncorrelated noise. Finally, we note that dynamic filtering has a couple of limitations with respect to spatially correlated noise, such as band-pass-limited additive noise (see discussion). We did not study this issue in more depth, as it would go beyond the scope of the present paper.

**Fig 20.**
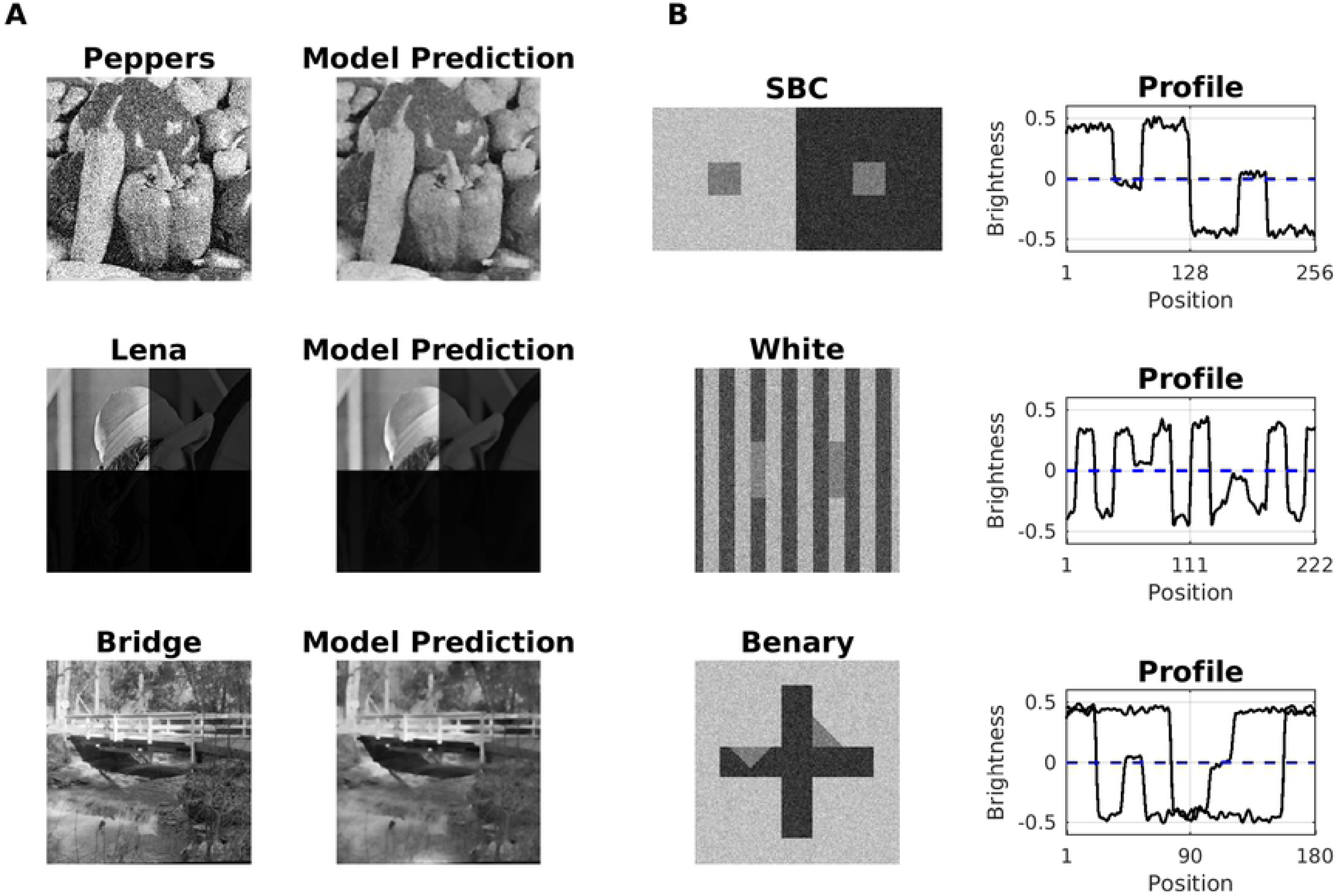
Real-world Image Processing. (A) Top: Peppers image with additive white noise (SNR=2.6266dB and PSNR=8.9813dB) along with the corresponding model output (SNR=5.4682db and PSNR=11.8228dB). The correlation coefficient between output and the noise-free original image was 0.9436. Middle: A high-dynamic-range version of the Lena image (where dynamic range of each quadrant decreases clockwise by one order of magnitude) and model output; the dynamic range of the input is 1, and that of the output is 0.9596. Bottom: Bridge image alongside with corresponding model output (SNR = 6.1084dB and PSNR = 13.6201dB). (B) Top: Simultaneous Brightness Contrast display with additive white noise and corresponding brightness profile (black line) as predicted by the model. The dashed line indicates the gray level of the gray squares. Middle: White Effect. Bottom: Benary Cross.

## Discussion

The perceived luminance (brightness) of target structure is highly sensitive to its spatial context. Despite of many modeling attempts for brightness, we still have not arrived at a detailed understanding of the corresponding neuronal information processing principles. With our model, we emphasize the role of predictive coding in brightness perception. Predictive coding for us simply means “what can be predicted will be suppressed”, and our dynamic filtering aims at reducing redundancy in boundary maps. Coding strategies that aim at reducing activity and thus energy expenditure in organisms are consistent, for example, with efficient coding [35–37, 90, 91], predictive coding [38], whitening [90] or response equalization [40]. In this sense, we propose that brightness perception is the consequence of suppressing redundant (i.e., predictable) information. Our model is build on the latter idea(s), and apart from being able to process real-world images, it predicts a bigger set of visual illusions than any other previously published model.

Our focus is thereby on low-level vision, as our model simulates the activity of simple and complex cells of the primary visual cortex. For each input, the model learns a filter kernel by identifying redundant patterns in (simulated) complex cell responses (i.e., the edge map), and subsequently uses the filter kernel to suppress redundant information (dynamic filtering). Dynamic filtering amounts to response equalization of simulated complex cell responses, much like the previously proposed “Whitening-by-Diffusion” method which directly acts on the (Fourier) amplitude spectrum [40]. The equalized responses are subsequently used for creating a representation of the sensory input by filling-in (brightness estimation). Nevertheless is dynamic filtering a global mechanism, which was adopted for the ease of implementation. In the primary visual cortex, we expect that dynamic filtering acts in a more local fashion, but still on a spatial scale that exceeds the typical receptive field sizes of V1 neurons. Such non-local mechanisms could be biologically implemented by feedback from mid-level visual neurons with sufficiently big receptive fields for detecting non-local correlations in activity.

We believe that our success in predicting a relatively large number of visual illusions lends some support to our proposed computational principle. Without changing any of our model’s parameter values, we are able to predict Simultaneous Brightness Contrast (SBC), White’s Effect, Reverse Contrast, Benary’s Cross, Todorovic’s illusion (with variations), the Dungeon Illusion, the Checkerboard Illusion, Shevell’s Ring, the Craik-O’Brien-Cornsweet effect (COCE), the Hermann/Hering grid, the corrugated grid, Chevreul’s illusion (including the luminance pyramid), Grating Induction (GI), and Mach Bands. Additionally, for some of the illusions, we were able to replicate data from psychophysical experiments (SBC, White, Reverse Contrast, Hermann/Hering grid, Todorovic, GI, and Mach Bands). By all the cheers we must not forget to describe some of the limitations of our model. We cannot predict illusions – without modifying the current parameters – such as achromatic neon “color” spreading, the Ehrenstein illusion, Chubb’s Illusion and some variations of the Hermann/Hering grid; Reverse Contrast with different grouping factors, SBC with articulated noise, and Mach Bands with an adjacently placed stimuli.

A further limitation is handling visual illusions where a target patch is surrounded by an articulation pattern instead of homogeneous luminance. Articulation patterns would introduce additional spatial redundancy into a luminance display, and this excess redundancy would be eventually learned by the kernel for dynamic filtering. As a consequence, dynamic filtering may modify the target’s edge representation in an unpredictable way. For brightness estimation, this could mean, for instance, that assimilation effects turn into contrast effects or vice versa. This behavior appears to be inconsistent with current psychophysical observations [92], and may hint at additional mechanisms that need to be considered. It cannot be ruled out that additional mechanisms reduce the redundancy along other stimulus dimensions as well, for example luminance, relative contrast, or auto-correlation. The global nature of the dynamic filter represents another trade-off. In order to learn the kernel for dynamic filtering, we sample patches randomly across the input. As a consequence, local statistical information between (unrelated) patches that are far away from each other could be intertwined, affecting their brightness predictions. A possible solution would be to introduce local constrains upon sampling, or as well to introduce a local normalization function, which takes into account local spatial auto-correlations.

### Comparison with other models that predict contrast and assimilation

Our predictive coding approach produces contrast effects by enhancing non-redundant edges through dynamic filtering according to Scenario 1. Assimilation effects are generated by suppressing redundant edges as a function of their relative intensity with respect to the other edges (Scenario 2). We thus do not make any explicit assumption about how a visual target is related to its context in terms of segmentation, or belongingness and perceptual frameworks, respectively. We do not require the categorization of image features either. In this sense, our predictive coding approach (as implemented by dynamic filtering) is more general than previous computational proposals and theories [4, 22, 28], which purport that a stimulus is divided into perceptual frameworks based on anchors [4] or T-junctions [22]. However, it is not clear whether anchors or T-junctions are sufficiently robust cues in real-world images, and actually few previously published models demonstrated the processing of real-world images.

Dynamic filtering is sensitive to the correlation structure of spatial patterns in order to generate contrast and assimilation effects. In this way, the output of our model would not be significantly affected if uncorrelated noise was added to the input. Yet multi-scale models are highly sensitive to additive noise [9–11], because their predictions depend on a careful re-adjustment of filter responses according to the contrast-sensitivity function [8]. Thus, if noise was added to contrast and assimilation displays, then corresponding predictions would be altered, because of corresponding changes in the spatial frequency spectrum [12].

Our model adapts to the statistical structure of each input image. This is to say that we do not evaluate each input image in a previously learned long-term statistical context. A long-term statistical context usually is learned from a big number of input samples in order to derive feature-specific probability distributions. In connection with brightness, a relationship between occurrence frequency of certain types of natural images and brightness perception has been proposed [31–33]. The main limitation of such models is that they require an enormous amount of data, and that visual illusions act much like an associative trigger or they are perceived according to humdrum Bayesian inference.

## Conclusion

In conclusion, this study provides a proof of concept of a hypothetical information processing strategy for visual system, based on economizing edge representations. Our predictions are reliant on the self-structure of the visual input, but not on accumulated visual experience, spatial frequency representations, or predefined detectors. Our proposed mechanism does not exclude information processing principles like accumulating visual experience or spatial frequency representations, and should be considered as being complementary to these. Finally, future work should address the understanding of how the statistical structure of the context surrounding a target patch influences its appearance. We also plan to study how different noise structures (as narrow-band, oriented, or correlated) influences the predictions of our model. Our predictive coding hypothesis should be compatible with all levels of information processing. This means that redundancy reduction likely might apply to higher-order patterns and shapes that form the primitives for object recognition.

## S1 Appendix. Gabor filters

Mathematically, our set of Gabor filters is parameterized as:

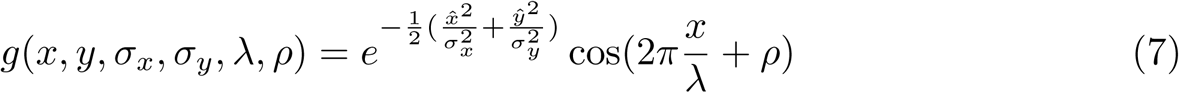

*ρ* is the phase, the parameters (*x, y*) are the spatial coordinates; *σ*_*x*_ and *σ*_*y*_ are the standard deviation along *x* and *y*, respectively; *λ* indicates the wavelength; and *θ* the orientation. We used *λ ∈* {4, 8}, eight orientations 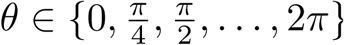, and Gabor filters with even and odd symmetry 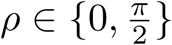. The standard deviations *σ*_*x*_ and *σ*_*y*_ depend on the wavelength and bandwidth *bw* = 2 as 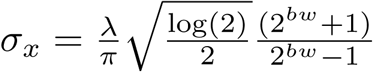 and 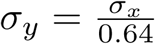.

The ON and OFF sub-fields of a Gabor filter g are normalized independently via:

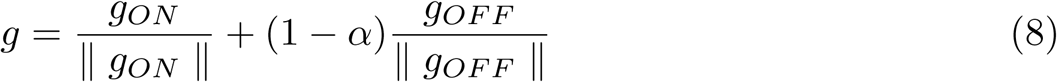

where ∥ ∥indicates Frobenius norm, and indicate *g*_*ON*_ and *g*_*OFF*_ region, respectively, of filter *g*. The parameter allows to control the sensitivity to luminance and is fixed at 0.1 (note that for *α* = 0 the neuron would not respond to homogeneous regions of luminance).

## S2 Appendix. Energy Map

Firstly, the complex cells were computed directly from the activity of the contrast-channel using the local energy model [42, 43, 51, 52] defined as:

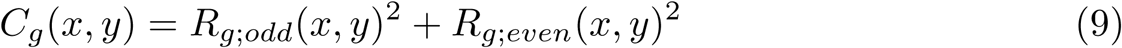

where *R*_*g,odd*_ and *R*_*g,even*_ indicates a pair of Gabor filters with identical orientation and spatial frequency, but different phase (*R*_*g,even*_ with *ρ* = 0, and *R*_*g,odd*_ with *ρ* = *π/*2).

Finally, the local energy map E was computed as:

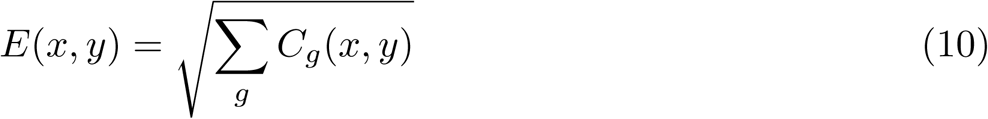

## S3 Appendix Dynamic filtering with zero-phase whitening (ZCA)

Initially, we sub-sampled the energy map E to half of its original size:

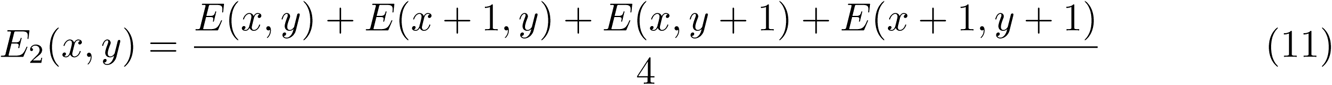

where *x, y* ∈ {1, 3, 5,… *n* − 1} are spatial indices. Because ZCA decorrelates intensity variations at the pixel level, the two-fold reduction in spatial scale has three advantages. First, the computational cost of ZCA is reduced. Second, the sensitivity to high spatial frequencies (and thus noise) is reduced. Third, because the edges on the energy map typically span more than one pixel in width, the scale reduction retained the intensity variations between different edges, while in turn reduced intensity variation along the edges. This led to less variable and thus improved contour map, what in turn facilitates the decorrelation between edges when applying the ZCA method.

Next, a set of 10000 patches of size 17 *×* 17 pixels was extracted randomly from the sub-sampled energy map *E*_2_. To be computationally tractable the set was normalized (extracting the mean and dividing by deviation) and cast into a matrix *X* of dimension 172 *×* 10000 such that the columns of the matrix represents each patch.

The zero-phase whitening (ZCA) transformation [55] consists in finding a symmetrical matrix *W* of dimension 17^2^ *×* 17^2^ such that - after applying it to *X* - the spatial correlations between the patches are eliminated (i.e., the covariance matrix after transformation is equal to the identity matrix). Then, the columns of *W* form a base in which the patches are decorrelated. Because *W* is symmetric, the columns of *W* are identical up to cyclic change in rows (i.e., the columns of *W* are only differ by shifting their values cyclically). This property permits to select any column of *W*, center it, and reshape it to build the kernel for the dynamic filter (see further down). In order to compute *W* we can solve the linear equation:

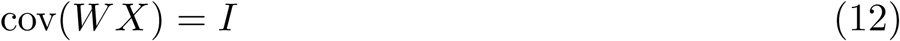

where *I* represents the identity matrix and cov indicates the covariance matrix. Due to *W* being symmetrical (i.e., *W* ^*T*^ = *W*), manipulating linear algebra operations, the last equation can be solved by

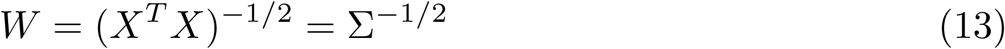

where Σ = cov(*X*) (i.e., represents the covariance matrix of *X*).

However, Eq 13 is very sensitive to high frequencies and isolated points (usually noise). This issue can be alleviated by introducing a regularization parameter and / or compression. We thus express the covariance matrix Σ by singular value decomposition Σ = *USV* ^*T*^ (using Matlab’s svd function) and add a regularization parameter according to Σ = *U* (*S* + *ϵI*)*V* ^*T*^, where *c* was set to 0.01 * max(*S*) and *I* is the identity matrix, and *S* is the matrix with singular values along its diagonal. In addition we reduce the dimension of the singular matrix *S* such that only the highest 50 singular values were kept and the remaining values were set to zero. The compressed matrix is 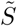, and we have

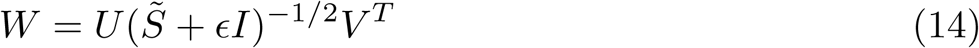

By imposing symmetry (*V* is substituted by *U*) we arrive at

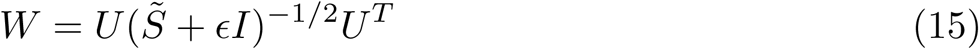

We define the dynamic filter *F* as the column of *W* with index 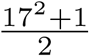 what corresponds to the centered column relative to the patches. The dynamic filter is extended (using Matlab’s imresize) and reshaped (using Matlab’s reshape) into a 2-dimensional matrix with size (2 *×* 17 − 1) *×* (2 *×* 17 − 1). Finally, the filter was normalized as 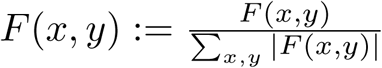 to be computationally tractable.

## S4 Appendix Solving Eq 5

The following gradient descent method served to compute the minima of the objective function *E*(*z*) of Eq 5. The iterative method is formulated as:

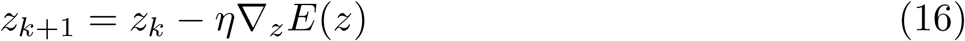

where *η* = 0.1 is a learning parameter. Because the objective function involves a norm and a linear operator (convolution), we computed the gradient ∇_*z*_*E*, indirectly, by using the definition of the directional derivative as:

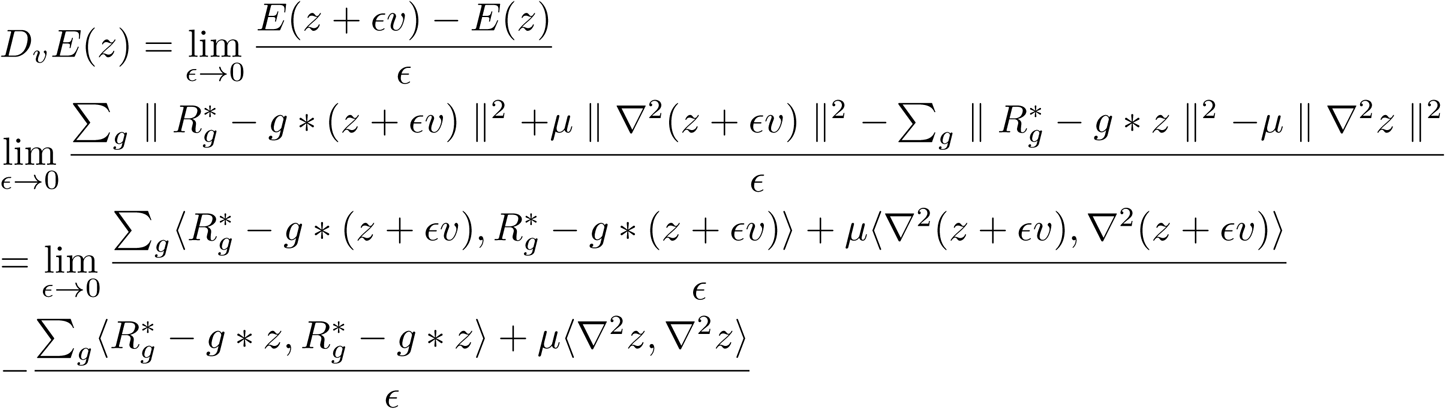

with the scalar product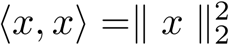. A straightforward manipulation of the scalar products and applying the limit we obtain:

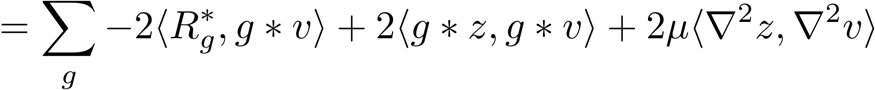

Now, isolating the directional vector v in the scalar product

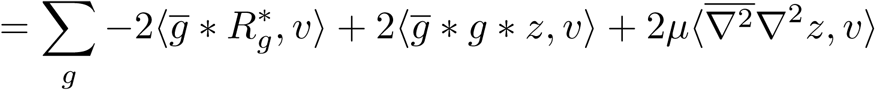

where filters with a bar indicate the corresponding complex adjoint. In particular, it holds:

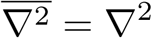

whereas taking the adjoint of Gabor filter g correspond to the same Gabor filters (as defined above in S1 Appendix) but with opposite sign in the phase parameter i.e.,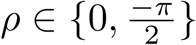. Finally using the the identity *D*_*v*_*E*(*z*) = ⟨∇_*z*_*E*_(_*z*), *v*⟩ for the directional derivative, we obtain:

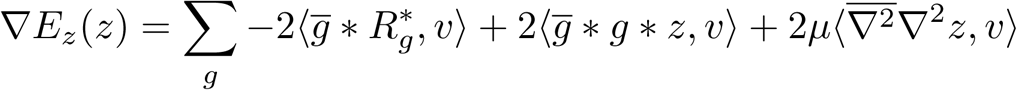

## Acknowledgments

We thank the Visca Group for laboratory space, and GonÇalo Correia for useful discussions. Funding for this project was provided by the Ministry of Economy and Competitiveness (Spain). Project numbers: PSI2010-18139-P, PSI2013-41568-P, PSI2014-57454-P, PGC2018-099506-B-I00 and PGC2018-096074-B-100.

## References

1. Rossi, A. F., Paradiso, M. A. Neural correlates of perceived brightness in the retina, lateral geniculate nucleus, and striate cortex. Journal of Neuroscience 1999, 19(14), 6145–6156. https://doi.org/10.1523/jneurosci.19-14-06145.1999

2. Barkan Y, Spitzer H, Einav S. Brightness contrast-contrast induction model predicts assimilation and inverted assimilation effects. Journal of Vision 2008;8(7):27. doi: https://doi.org/10.1167/8.7.27.

3. White M. The effect of the nature of the surround on the perceived lightness of grey bars within square wave test gratings. Perception. 1981; 10, no. 2 215–30. doi:10.1068/p100215.

4. Gilchrist A, Kossyfidis C, Agostini T, Li X, Bonato F, Cataliotti J, et al. An anchoring theory of lightness perception. Psychological Review. 1999. 106(4), 795–834. https://doi.org/10.1037/0033-295X.106.4.795

5. Anderson BL, Whitbread M, de Silva C. Lightness, brightness, and anchoring. Journal of Vision 2014, Aug 2;14(9):7–7.

6. Maniatis LM. A theory divided: Current representations of the anchoring theory of lightness contradict the original’s core claims. Vol. 102, Vision Research. Elsevier Ltd; 2014. p. 89–92.

7. Hong S, Grossberg S. A neuromorphic model for achromatic and chromatic surface representation of natural images. Neural Networks. 2004; Volume 17, Issues 5–6, Pages 787–808, ISSN 0893-6080, https://doi.org/10.1016/j.neunet.2004.02.007.

8. DeValois RL, DeValois KK. Spatial Vision. Annual Review of Psychology. 1988. 31(1) 309–341.

9. Blakeslee B, McCourt ME. A multiscale spatial filtering account of the White effect, simultaneous brightness contrast and grating induction. Vision Research. 1999; Volume 39, Issue 26, Pages 4361–4377, ISSN 0042-6989, https://doi.org/10.1016/S0042-6989(99)00119-4.

10. Blakeslee B, McCourt ME. A unified theory of brightness contrast and assimilation incorporating oriented multiscale spatial filtering and contrast normalization. Vision Research. 2004; Volume 44, Issue 21, Pages 2483–2503, ISSN 0042-6989, https://doi.org/10.1016/j.visres.2004.05.015.

11. Otazu X, Vanrell M, Alejandro Párraga C. Multiresolution wavelet framework models brightness induction effects. Vision Research. 2008, Volume 48, Issue 5, Pages 733–751, ISSN 0042-6989, https://doi.org/10.1016/j.visres.2007.12.008.

12. Betz T, Shapley R, Wichmann FA, Maertens M. Noise masking of White’s illusion exposes the weakness of current spatial filtering models of lightness perception. Journal of Vision. 2015; 15(14):1. http://jov.arvojournals.org/article.aspx?doi=10.1167/15.14.1

13. Horn BKP. Determining lightness from an image. Comput Graph Image Process. 1974;

14. Land EH, McCann JJ. Lightness and retinex theory. J Opt Soc Am. 1971;

15. Gerrits HJM, Vendrik AJH. Simultaneous contrast, filling-in process and information processing in man’s visual system. Exp Brain Res. 1970;

16. Cohen MA, Grossberg S. Neural dynamics of brightness perception: Features, boundaries, diffusion, and resonance. Adv Psychol. 1987;

17. Keil MS, Cristóbal G, Hansen T, Neumann H. Recovering real-world images from single-scale boundaries with a novel filling-in architecture. Neural Networks. 2005;

18. Grossberg S, Todorovic D. Neural dynamics of 1-D and 2-D brightness perception: A unified model of classical and recent phenomena. Percept Psychophys. 1988 May;43(3):241–77.

19. Pessoa L, Thompson E, Noe A. Finding out about filling-in: A guide to perceptual completion for visual science and the philosophy of perception. Behav Brain Sci. 1998;21(6):723–802.

20. Komatsu H. The neural mechanisms of perceptual filling-in. Vol. 7, Nature Reviews Neuroscience. Nature Publishing Group; 2006. p. 220–31.

21. Grossberg S. 3-D vision and figure-ground separation by visual cortex. Perception & Psychophysics. 1994.

22. Ross WD, Pessoa L. Lightness from contrast: A selective integration model. Percept Psychophys. 2000;62(6):1160–81.

23. Yazdanbakhsh A, Arabzadeh E, Babadi B, Fazl A. Munker-White-like illusions without T-junctions. Perception. 2002;31(6):711–5.

24. Howe PDL. White’s effect: Removing the junctions but preserving the strength of the illusion. Perception. 2005;

25. Bressan P. Explaining lightness illusions. Perception. 2001;

26. Todorović D. Lightness and junctions. Perception. 1997;

27. Keil MS, Cristóbal G, Neumann H. Gradient representation and perception in the early visual system-A novel account of Mach band formation. Vision Res [Internet]. 2006 Sep [cited 2020 Mar 17];46(17):2659–74. Available from: http://www.ncbi.nlm.nih.gov/pubmed/16603218

28. Domijan D. A Neurocomputational account of the role of contour facilitation in brightness perception. Front Hum Neurosci. 2015;9:93. Available from: http://www.ncbi.nlm.nih.gov/pubmed/25745396

29. Keil MS. Local to global normalization dynamic by nonlinear local interactions. Phys D Nonlinear Phenom. 2008;

30. Neumann H. Mechanisms of neural architecture for visual contrast and brightness perception. Neural Networks. 1996;

31. Yang Z, Purves D. The statistical structure of natural light patterns determines perceived light intensity. Proc Natl Acad Sci. 2004 Jun 8;101(23):8745–50. Available from: https://www.pnas.org/content/101/23/8745

32. Purves D, Williams SM, Nundy S, Lotto RB. Perceiving the Intensity of Light. Psychol Rev. 2004 Jan; 111(1):142–58. Available from: http://www.ncbi.nlm.nih.gov/pubmed/14756591

33. Corney D, Lotto RB. What Are Lightness Illusions and Why Do We See Them? PLoS Comput Biol. 2007;3(9):e180. Available from: http://dx.plos.org/10.1371/journal.pcbi.0030180

34. Morgenstern Y, Rukmini D V., Monson BB, Purves D. Properties of artificial neurons that report lightness based on accumulated experience with luminance. Front Comput Neurosci. 2014;

35. Vincent BT, Baddeley RJ, Troscianko T, Gilchrist ID. Is the early visual system optimised to be energy efficient? In: Network: Computation in Neural Systems. 2005; 16. 175-90. 10.1080/09548980500290047.

36. Levy WB, Baxter RA. Energy Efficient Neural Codes. Neural Comput. 1996;

37. Barlow HB. Possible principles underlying the transformation of sensory messages. Sens Commun. 1961; Available from: https://ci.nii.ac.jp/naid/10012745911/

38. Srinivasan M V., Laughlin SB, Dubs A. Predictive coding: A fresh view of inhibition in the retina. Proc R Soc London - Biol Sci. 1982;

39. Atick JJ, Redlich AN. What Does the Retina Know about Natural Scenes? Neural Comput. 1992;

40. Keil MS. Does face image statistics predict a preferred spatial frequency for human face processing? Proc R Soc B Biol Sci. 2008;

41. Jones JP, Palmer LA. An evaluation of the two-dimensional Gabor filter model of simple receptive fields in cat striate cortex. J Neurophysiol. 1987;

42. Morrone MC, Owens RA. Feature detection from local energy. Pattern Recognit Lett. 1987;

43. Morrone MC, Burr DC. Feature detection in human vision: a phase-dependent energy model. Proc R Soc Lond B Biol Sci. 1988;

44. Hubel DH, Wiesel TN. Receptive fields of single neurones in the cat’s striate cortex. J Physiol. 1959;

45. Daugman JG. Uncertainty relation for resolution in space, spatial frequency, and orientation optimized by two-dimensional visual cortical filters. J Opt Soc Am A. 1985;

46. Jones JP, Palmer LA. An evaluation of the two-dimensional Gabor filter model of simple receptive fields in cat striate cortex. J Neurophysiol. 1987;

47. Marcelja S. Mathematical description of the responses of simple cortical cells. J Opt Soc Am. 1980;

48. Komatsu H, Murakami I, Kinoshita M. Surface representation in the visual system. Cognitive Brain Research. 1996;

49. Rossi AF, Rittenhouse CD, Paradiso MA. The representation of brightness in primary visual cortex. Science (80-). 1996;

50. Dai J, Wang Y. Representation of surface luminance and contrast in primary visual cortex. Cereb Cortex. 2012;

51. Adelson EH, Bergen JR. Spatiotemporal energy models for the perception of motion. J Opt Soc Am A. 1985;

52. Pollen DA, Ronner SF. Visual Cortical Neurons as Localized Spatial Frequency Filters. IEEE Trans Syst Man Cybern. 1983;

53. Hubel DH, Wiesel TN. Receptive fields, binocular interaction and functional architecture in the cat’s visual cortex. J Physiol. 1962;

54. De Valois RL, Albrecht DG, Thorell LG. Spatial frequency selectivity of cells in macaque visual cortex. Vision Res. 1982;22(5):545–59.

55. Bell AJ, Sejnowski TJ. The “independent components” of natural scenes are edge filters. Vision Res. 1997;

56. Khintchine A. Korrelationstheorie der stationären stochastischen Prozesse. Math Ann. 1934; Dec, 109(1):604–15. Available from: http://link.springer.com/10.1007/BF01449156

57. Wiener N. Generalized harmonic analysis. Acta Math. 1930;55(0):117–258. Available from: http://projecteuclid.org/euclid.acta/1485887877

58. Grossberg S, Hong S. A neural model of surface perception: Lightness, anchoring, and filling-in. Spat Vis. 2006;

59. Logvinenko AD, Kane J. Hering’s and Helmholtz’s types of simultaneous lightness contrast. J Vis. 2005;

60. Blakeslee B, McCourt ME. Similar mechanisms underlie simultaneous brightness contrast and grating induction. Vision Res. 1997;

61. Diamond AL. Foveal simultaneous brightness contrast as a function of inducing, and test-field luminances. J Exp Psychol. 1953;

62. Kitterle FL. The effects of simultaneous and successive contrast on perceived brightness. Vision Res. 1972;

63. Stevens JC. Brightness inhibition re size of surround. Percept Psychophys. 1967;

64. William Yund E, Armington JC. Color and brightness contrast effects as a function of spatial variables. Vision Res. 1975;

65. Shi V, Cui J, Troncoso XG, Macknik SL, Martinez-Conde S. Effect of stimulus width on simultaneous contrast. PeerJ. 2013;

66. Benary W. Beobachtungen zu einem Experiment über Helligkeitskontrast. Psychol Forsch. 1924;

67. Salmela VR, Laurinen PI. Low-level features determine brightness in White’s and Benary’s illusions. Vision Res. 2009 Apr 29; 49(7):682–90. Available from: http://www.ncbi.nlm.nih.gov/pubmed/19200439

68. Economou E, Zdravković S, Gilchrist A. Grouping Factors and the Reverse Contrast Illusion. Perception. 2015;

69. Taya R, Ehrenstein WH, Cavonius CR. Varying the strength of the Munker-White effect by stereoscopic viewing. Perception. 1995;

70. Ripamonti C, Gerbino W. Classical and inverted White’s effects. Perception. 2001;

71. Kingdom F, Moulden B. White’s effect and assimilation. Vision Res. 1991;31(1):151–9. Available from: http://www.ncbi.nlm.nih.gov/pubmed/2006548

72. Betz T, Shapley R, Wichmann FA, Maertens M. Testing the role of luminance edges in White’s illusion with contour adaptation. J Vis. 2015 Aug 1;15(11):14. Available from: http://www.ncbi.nlm.nih.gov/pubmed/26305862

73. Güçlü B, Farell B. Influence of target size and luminance on the White-Todorović effect. Vision Res. 2005;

74. Hong SW, Shevell SK. Brightness contrast and assimilation from patterned inducing backgrounds. Vision Res. 2004; Volume 44, Issue 1, Pages 35–43, ISSN 0042-6989, https://doi.org/10.1016/j.visres.2003.07.010.

75. Baumgartner G. Indirekte Größenbestimmung der rezeptiven Felder der Retina beim Menschen mittels der Hermannschen Gittertäuschung. Pflugers Arch Gesamte Physiol Menschen Tiere. 1960;

76. Schiller PH, Carvey CE. The Hermann grid illusion revisited. Perception. 2005.

77. Keil MS. Smooth Gradient Representations as a Unifying Account of Chevreul’s Illusion, Mach Bands, and a Variant of the Ehrenstein Disk. Neural Comput. 2006 Feb 12;18(4):871–903.

78. Keil MS. Gradient representations and the perception of luminosity. Vision Res. 2007 Dec 1;47(27):3360–72.

79. Jameson D, Hurvich LM, Ratliff F. Mach Bands: Quantitative Studies on Neural Networks in the Retina. Am J Psychol. 1966;

80. Ross J, Concetta Morrone M, Burr DC. The conditions under which Mach bands are visible. Vision Res. 1989 Jan 1;29(6):699–715. Available from: https://www.sciencedirect.com/science/article/pii/0042698989900333

81. Pessoa L. Mach bands: how many models are possible? Recent experimental findings and modeling attempts. Vision Res. 1996 Oct;36(19):3205–27. Available from: http://www.ncbi.nlm.nih.gov/pubmed/8917780

82. Ratliff F. Why mach bands are not seen at the edges of a step. Vision Res. 1984;

83. Kingdom FAA. Mach bands explained by response normalization. Front Hum Neurosci. 2014;

84. Ratliff F, Milkman N, Rennert N. Attenuation of Mach bands by adjacent stimuli. Proc Natl Acad Sci U S A. 1983 Jul;80(14):4554–8. Available from: http://www.ncbi.nlm.nih.gov/pubmed/6576350

85. McCourt ME. A spatial frequency dependent grating-induction effect. Vision Res. 1982;

86. Keil MS. From Neuronal Models to Neuronal Dynamics and Image Processing. Biologically inspired Computer Vision: Fundamentals and Applications. wiley; 2015. p. 221–44. Available from: http://arxiv.org/abs/1801.08585

87. Grossberg S. Cortical dynamics of three-dimensional form, color, and brightness perception: I. Monocular theory. Percept Psychophys. 1987 Mar;41(2):87–116.

88. Keil MS. Neural architectures for unifying brightness perception and image processing [dissetation] Universität Ulm; 2003. Available from: https://www.researchgate.net/publication/35660056 Neural architectures for unifying brightness perception and image processing

89. Lerer A, Supér H, Keil MS. Luminance gradients and non-gradients as a cue for distinguishing reflectance and illumination in achromatic images: A computational approach. Neural Networks. 2019;110.

90. Atick JJ. Could information theory provide an ecological theory of sensory processing? Network: Computation in Neural Systems. 1992.

91. Simoncelli EP, Olshausen BA. Natural Image Statistics and Neural Representation. Annu Rev Neurosci. 2001;

92. Bressan P, Actis-Grosso R. Simultaneous Lightness Contrast on Plain and Articulated Surrounds. Perception. 2006 Apr 25;35(4):445–52. Available from: http://journals.sagepub.com/doi/10.1068/p5247

